# NMR reveals specific remodelling of protein folding landscapes in ionic liquids

**DOI:** 10.64898/2026.01.21.700775

**Authors:** Micael S. Silva, Aldino Viegas, Philip O’Toole, Sara S. Felix, Angelo Miguel Figueiredo, Eurico J. Cabrita

## Abstract

Ionic liquids (ILs) are designer solvents with tunable properties that have emerged as powerful cosolutes in protein science and biotechnology. While ILs are known to stabilize or destabilize proteins, their molecular mechanisms remain poorly defined, particularly regarding effects on the unfolded ensemble. Here, we use the metastable SH3 domain of the Drosophila adaptor protein Drk, which coexists in a slow two-state equilibrium between folded and unfolded states, as a model to dissect how ILs modulate protein conformational equilibria. Using nuclear magnetic resonance (NMR) spectroscopy, we show that cholinium glutamate ([Ch][Glu]) stabilizes the folded state by preferential exclusion from the protein surface, raising the barrier to unfolding in a manner reminiscent of osmolytes and crowding agents. In contrast, 1-butyl-3-methylimidazolium dicyanamide ([Bmim][dca]) shifts the equilibrium toward the unfolded state, not by destabilizing the native fold, but by stabilizing a compact, non-native α-helical conformation within the unfolded ensemble. Notably, this concentration-dependent mechanism differs from traditional denaturants such as urea or guanidinium chloride, which promote random-coil unfolded states. Kinetic measurements further reveal that [Ch][Glu] slows unfolding, whereas [Bmim][dca] slows folding, indicating that the two ILs reshape the folding landscape in opposite directions. These findings challenge the prevailing view that cosolutes act primarily on the folded state and establish unfolded-state stabilization as a critical determinant of protein stability. More broadly, this work provides a molecular framework for understanding how ILs reshape protein folding landscapes and offers design principles for tailoring ILs as stabilizers or destabilizers in biotechnology and therapeutic applications.

**Significance Statement:** Proteins exist in a delicate balance between folded and unfolded states, and cosolutes such as salts are generally viewed as acting only on the folded state. Here, we show that ionic liquids (ILs) are able to reshape this balance and uniquely modulate both folded and unfolded protein states through distinct molecular mechanisms. These results challenge the conventional view of protein stabilization/destabilization and demonstrate that unfolded-state modulation is a critical determinant of protein behavior. By establishing principles for how ILs alter protein folding landscapes, this work provides a framework for designing ILs with tailored effects, with implications for biocatalysis, protein engineering, and therapeutic formulation.

**Graphical Abstract:** 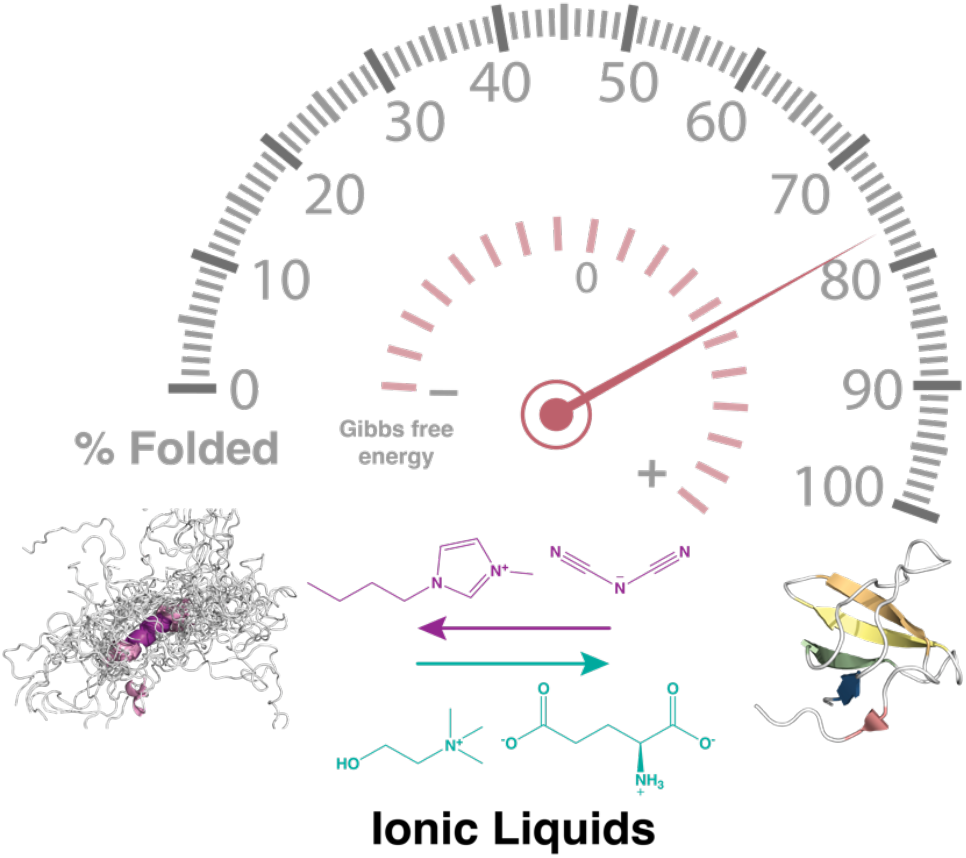

## Introduction

Since the pioneering work of Hofmeister over 130 years ago (1), ion-specific effects have been recognized as key modulators of protein properties. The Hofmeister series, originally developed to rank anions and cations by their efficiency in precipitating proteins, was later correlated with their ability to stabilize or destabilize proteins (2). Beyond proteins, Hofmeister effects have since been implicated in colloid, polymer, and surface chemistry (3).

In the past two decades, the development of tunable low-melting-point salts, known as ionic liquids (ILs), has greatly expanded the chemical space for exploring ion–protein interactions (4, 5). ILs, composed of organic cations paired with organic or inorganic anions, can form more than 10^6^ possible combinations with tunable polarity, hydrophobicity, and hydrogen-bonding capacity (6). Their versatility has led to broad applications, from biocatalysis (7) to protein stabilization (8). Both in neat and aqueous solutions, ILs can shift the delicate balance between folded and unfolded protein states, acting either as denaturants or stabilizers (9–11). Among the most studied are cholinium- and imidazolium-based ILs, which represent opposite ends of the biocompatibility spectrum (12, 13).

Empirical trends suggest that IL effects often follow the Hofmeister series, with anions exerting stronger influences than cations (14, 15). Yet, these effects are highly concentration-dependent and nonlinear, as the same ion can stabilize or destabilize a protein depending on mole fraction (16). At the molecular level, IL–protein interactions emerge from electrostatics, hydrogen bonding, dispersion, and excluded-volume effects, modulated by ion and protein hydration (17). Importantly, most studies focus on the folded ensemble, assuming minimal structural changes in unfolded states. However, evidence from intrinsically disordered proteins and molten globules demonstrates that unfolded ensembles are sensitive to ionic strength, solvation, and electrostatics, and are far from random coils (18–20). Ignoring their response risks incomplete interpretations of protein stability.

High-resolution, site-specific approaches are therefore essential to dissect how ILs reshape protein conformational equilibria (21). Nuclear magnetic resonance (NMR) spectroscopy is uniquely suited to this challenge, as it can resolve both folded and unfolded states, quantify thermodynamic parameters, and map subtle cosolute–protein interactions at atomic detail.

Here, as a model system, we employ the 59-residue SH3 domain of the Drosophila adapter protein Drk, which exists in near-equal equilibrium between folded (F) and unfolded (U) states under mild aqueous conditions (22, 23). These states exchange slowly on the NMR timescale (*k*_ex_ ≈ 2 s^−1^ at pH 6.0, 293 K) and can be simultaneously observed (**Fig. 1**), with the indole NH of W36 serving as a population marker (24).

**Fig. 1.**
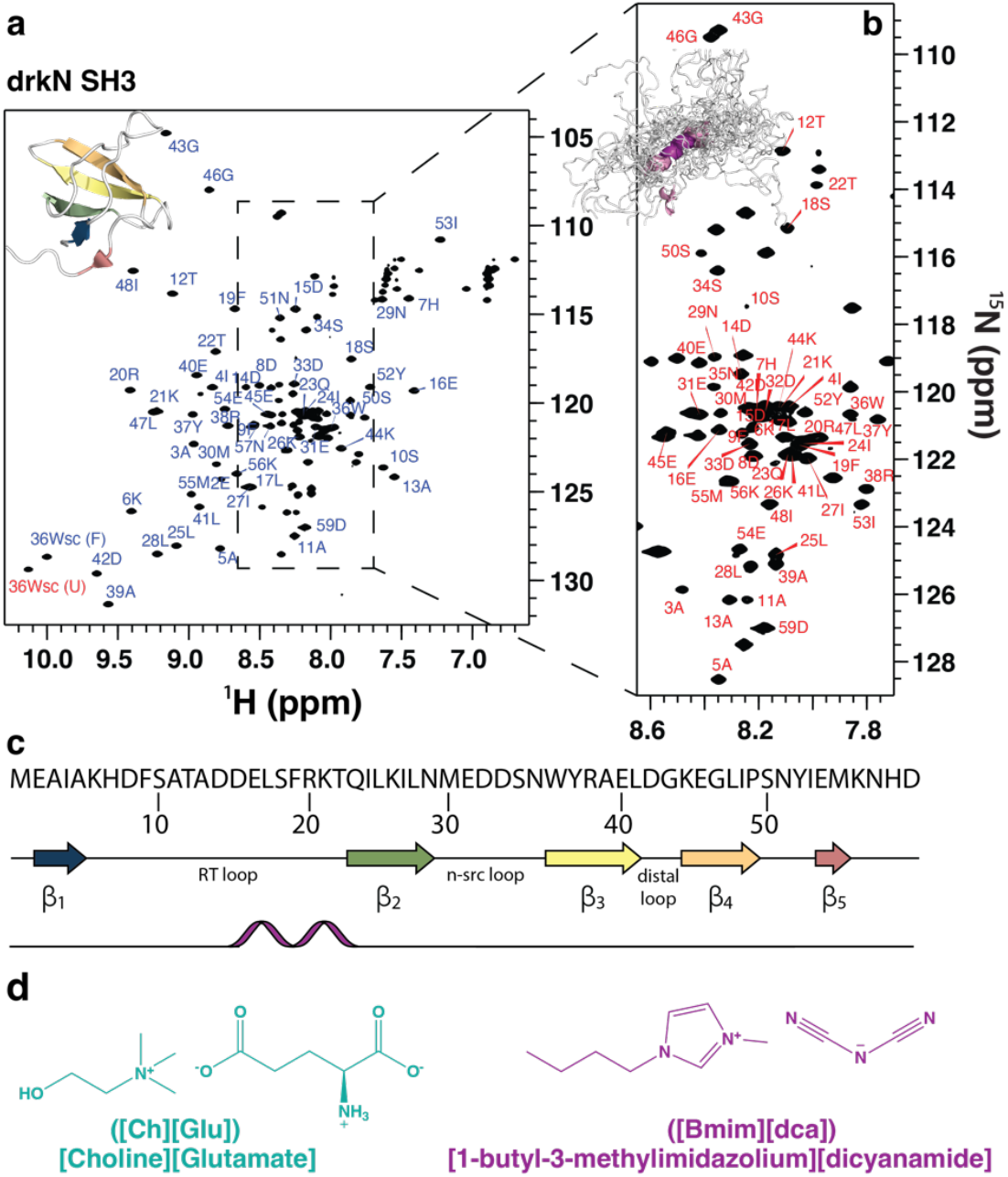
Folded and unfolded states of SH3 exist in slow exchange on the NMR chemical shift timescale. (**a, b**) [^1^H,^15^N]-HSQC spectra SH3 showing the assignment of amide NH peaks for the folded (a) and unfolded (b) states in water at 298 K, pH 7.1. The folded state is represented by a 3D structure (PDB: 2A36), and the unfolded state by 30 ensemble representatives (PED00022). (**c**) Primary sequence (top) and secondary structure of folded (middle) and unfolded (bottom) SH3. β-sheets are shown as arrows, loops as lines, and the α-helix in purple. The RT loop includes two irregular β-hairpins, and strands β4 and β5 are connected by a short 3_10_ helix. (**d**) Structures of the ionic liquids: choline glutamate ([Ch][Glu], left) and 1-butyl-3-methylimidazolium dicyanamide ([Bmim][dca], right).

Using this system, we investigate how two contrasting ILs – cholinium glutamate ([Ch][Glu]), a biocompatible stabilizer, and 1-butyl-3-methylimidazolium dicyanamide ([Bmim][dca]), a destabilizer – modulate the folding equilibrium of the SH3 domain. By integrating NMR structural analysis with thermodynamic and kinetic profiling, we reveal distinct molecular pathways: [Ch][Glu] stabilizes the folded state primarily through entropic excluded-volume effects, whereas [Bmim][dca] promotes unfolding by stabilizing a compact, non-native α-helical conformation. These findings advance our understanding of IL–protein interactions and provide a mechanistic framework for rational IL design in biotechnology.

## Results and Discussion

### IL-induced modulation of protein stability

To assess how ionic liquids (ILs) affect the folding equilibrium of the SH3 domain, we titrated the protein with increasing concentrations of [Ch][Glu] and [Bmim][dca] and monitored changes by [^1^H,^15^N]-HSQC spectroscopy (**Fig. S1a, S1b**). The intensity of the indole NH sidechain of W36, which is well resolved in both folded and unfolded states, was used as a proxy for conformational populations (**Fig. S1c, S1d**). Addition of [Ch][Glu] caused a progressive disappearance of the unfolded-state resonances, consistent with stabilization of the folded state (**Fig. S1a and c**). By contrast, [Bmim][dca] shifted the equilibrium in the opposite direction, favoring the unfolded ensemble (**Fig. S1b and d**).

Given the pronounced electrostatic surface potential of SH3 (pI ≈ 4.6), we next asked whether these effects could be explained by (i) nonspecific ionic strength (electrostatic shielding), (ii) specific ion-protein interactions, or (iii) a unique “ionic liquid effect” arising from ion pairing. To disentangle these contributions, we repeated the titrations with the individual ion components: [Ch]Cl, [Bmim]Cl, Na[Glu], and Na[dca], using NaCl as a reference (**Fig. S2**). Again, the folded (F) and unfolded (U) populations were quantified from W36 indole N^ε1^-H^ε1^ peak volumes (*V*_f_ and *V*_u_), which are proportional to their equilibrium populations (*p*_f_ and *p*_u_). The modified standard Gibbs free energy of unfolding was then calculated as:

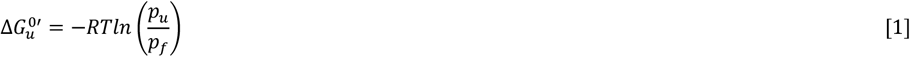

where *R* is the gas constant and *T* is the absolute temperature.

Assuming a two-state unfolding model (F ⇌ U), the dependence of Δ*G*^0’^_u_ on IL concentration can be described by:

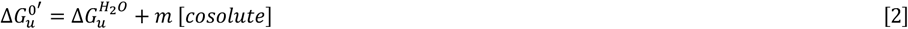

where Δ*G*^H2O^_u_ is the free energy of unfolding in water, and the slope (*m*-value) quantifies the cosolute’s effect: positive *m*-values indicate stabilization, while negative *m*-values indicate destabilization (25).

The extracted Δ*G*^0’^_u_ and *m*-values (**Fig. 2, Table 1**) reveal a striking contrast. [Ch][Glu] stabilizes SH3 with an *m*-value of +1.9 ± 0.1 kcal·mol^− 1^·M^−1^, whereas [Bmim][dca] strongly destabilizes it (– 7.7 ± 0.3 kcal·mol^−1^·M^−1^). None of the individual salts reproduced the magnitude of the corresponding ILs, indicating that the observed effects cannot be explained solely by ionic strength. Consistent with this, earlier NMR studies showed that even high NaCl concentrations fail to fully stabilize SH3 by electrostatic screening alone (26).

**Table 1.**
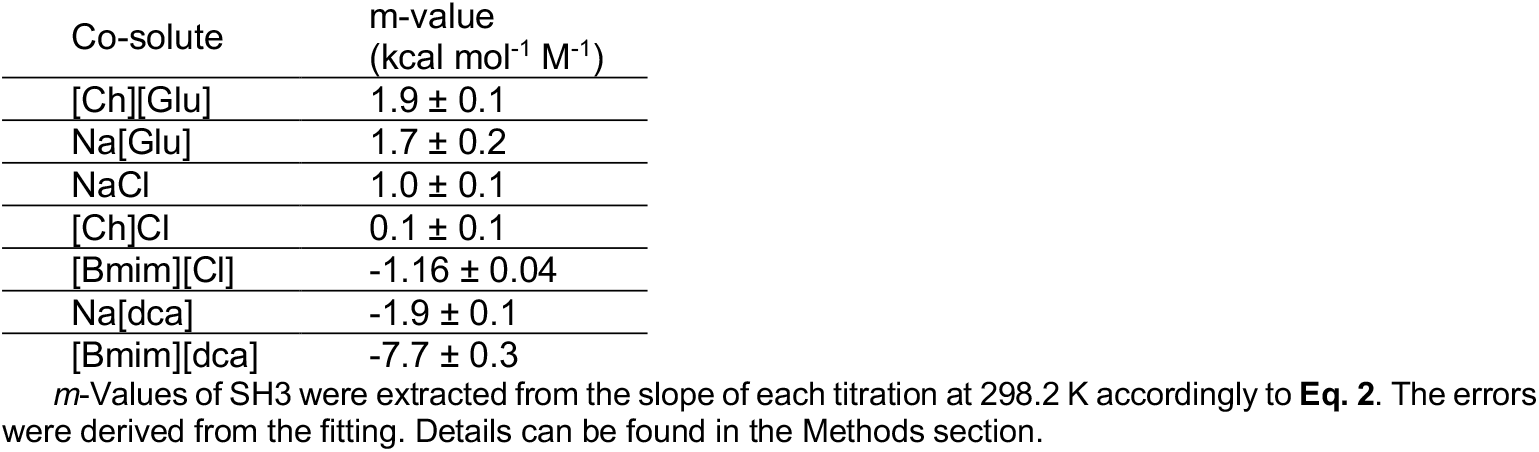
*m*-Values of SH3 in the presence of different cosolutes.

| Co-solute | $m$-value<br>(kcal mol<sup>-1</sup> M<sup>-1</sup>) |
| --- | --- |
| [Ch][Glu] | 1.9 ± 0.1 |
| Na[Glu] | 1.7 ± 0.2 |
| NaCl | 1.0 ± 0.1 |
| [Ch]Cl | 0.1 ± 0.1 |
| [Bmim][Cl] | -1.16 ± 0.04 |
| Na[dca] | -1.9 ± 0.1 |
| [Bmim][dca] | -7.7 ± 0.3 |

$m$ -Values of SH3 were extracted from the slope of each titration at 298.2 K accordingly to **Eq. 2**. The errors were derived from the fitting. Details can be found in the Methods section.

**Fig. 2.**
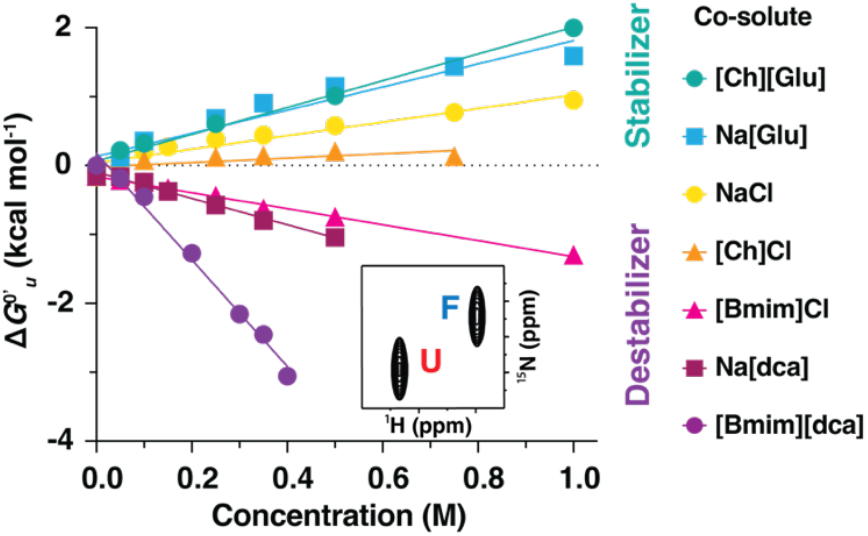
Cosolute-induced stabilization and destabilization of SH3. Effect of [Ch][Glu] and [Bmim][dca] and corresponding ionic salts on Δ*G*^*0’*^_u_ of SH3. In water, the populations are nearly equal (Δ*G*^0^_u_ ≈ 0, pf ≈ pu) (inlay). Δ*G*^0^_u_ and *m*-values were determined according to equations 1 and 2, respectively, with errors propagated from peak volume measurements.

For [Ch][Glu], stabilization is primarily driven by the glutamate anion (**Fig. 2, turquoise**). Na[Glu] alone produces nearly the same stabilizing effect (*m* = +1.7 ± 0.2), whereas [Ch]Cl is essentially neutral (*m* = +0.1 ± 0.1). This points to an overcompensating stabilizing effect of [Glu]^−^, consistent with its placement at the stabilizing end of the Hofmeister series (27). Similar effects have been reported for Na[Glu] and K[Glu] (28) and for other cholinium-based ILs with dicarboxylate anions, where hydrogen bonding and electrostatic interactions of the anion dominate (29).

In contrast, both components of [Bmim][dca] are destabilizing. Na[dca] alone decreases stability (*m* = –1.9 ± 0.1), and [Bmim]Cl has a comparable effect (*m* = –1.16 ± 0.04). These contributions likely reflect the hydrophobic, surface-active nature of the imidazolium cation and the high polarizability and low charge density of the [dca]^−^ anion, both of which promote interactions with nonpolar residues and lower protein–water interfacial tension (30). Strikingly, their combination in [Bmim][dca] produces a synergistic destabilization far stronger than the sum of the individual salts, revealing a genuine “ionic liquid effect.”

Overall, the tested cosolutes can be ranked by their impact on SH3 stability as follows:

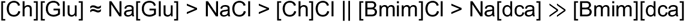

Where the double bar (||) indicates the transition from stabilizing to destabilizing behavior. The trends are broadly consistent with Hofmeister expectations (16): cholinium ILs are dominated by their stabilizing anions, whereas imidazolium ILs combine destabilizing cations and anions to act as potent denaturants. Nevertheless, the extreme destabilization by [Bmim][dca] highlights that IL effects cannot be reduced to ion identity alone but also depend on ion pairing, IL composition, concentration, protein surface topology, and specific ion–protein interactions (8, 13, 21).*m*-Values of SH3 were extracted from the slope of each titration at 298.2 K accordingly to **Eq. 2**. The errors were derived from the fitting. Details can be found in the Methods section.

### IL-Protein interactions

Having observed that different salts modulate SH3 stability and that simple electrostatic shielding cannot fully explain the effects, we investigated the mechanisms underlying protein (de)stabilization. Our previous studies (15, 31) showed that both electrostatic and hydrophobic interactions are critical for understanding IL effects on protein stability. To identify and map these interactions, we analyzed backbone amide chemical shift perturbations (CSPs) from [^1^H,^15^N]-HSQC spectra acquired in the presence of increasing concentrations of [Ch][Glu] and [Bmim][dca] (**Fig. S1**) and of the corresponding ionic species (**Fig. S2**). For clarity, **Fig. 3** shows representative conditions: 0.35 M [Ch][Glu] (78% folded, F[Ch][Glu]; 22% unfolded, U[Ch][Glu]) and 0.15 M [Bmim][dca] (14% folded, F_[Bmim][dca]_; 86% unfolded, U_[Bmim][dca]_). CSPs were calculated as combined chemical shift changes (Δδ_comb_; (32)) and mapped onto the folded structure (PDB: 2A36; (33)) and a representative structure from the unfolded ensemble (**Fig. 3**). Here, the chosen unfolded conformer reflects the experimental dataset in which residues 15–22 populate the α-region >70% of the time with ~30% α-helix within the ensemble (PED00022; (23, 34, 35)).

**Fig. 3.**
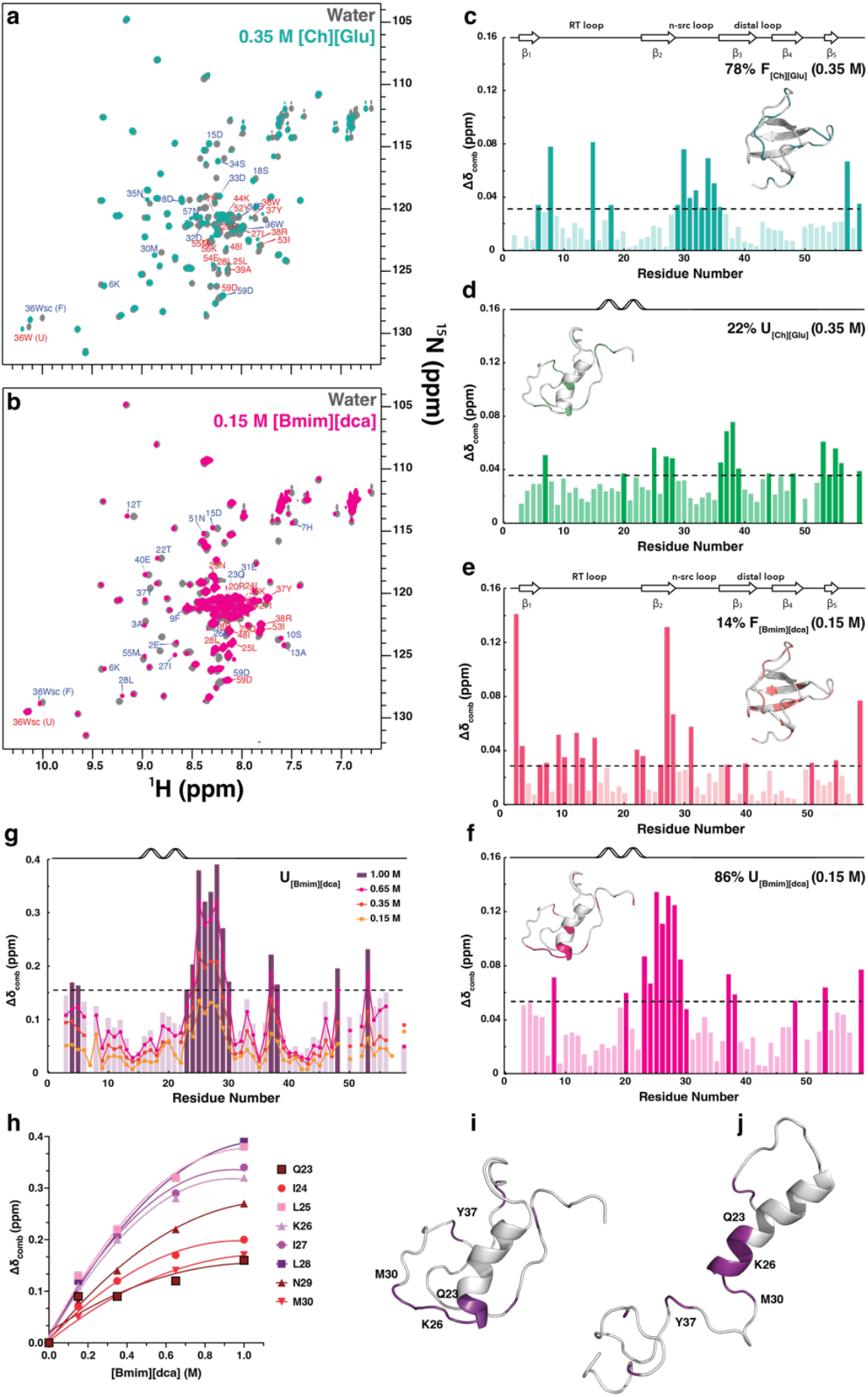
Ionic liquid–SH3 interactions. Overlay of 2D [^1^H,^15^N]-HSQC of SH3 in water (gray) and (**a**) 0.35 M [Ch][Glu] (turquoise) or (**b**) [Bmim][dca] (pink); labels shown for the most affected residues; **(c-d)** Combined chemical shift (Δδ_comb_) profiles for SH3 at 0.35 M of [Ch][Glu] in the folded (turquoise) and unfolded (green) states, respectively; residues above the threshold (dashed line) are colored darker. **(e-f)** Δδ_comb_ profiles at 0.15 M [Bmim][dca] for folded (salmon) and unfolded (pink) states; residues above threshold are colored darker. **(g)** Overlay of Δδ_comb_ profiles for unfolded SH3 at increasing [Bmim][dca]: 0.15 M (orange), 0.35 M (pink), 0.65 M (hot pink), and 1.0 M (dark purple). The dashed line marks the cutoff determined at 1.0 M. Above each plot is indicated the protein secondary structure. Inset structures map affected residues onto the folded state (PDB: 2A36) and a representative unfolded conformer (PED00022, #576). Δδ_comb_ values were referenced to F_water_ and U_water_. **(h)** Δδ_comb_ versus [Bmim][dca] for Q23–M30. **(i–j)** Representative unfolded structures from PED00022: **(i)** #576 and (**j**) #1195, with affected residues in purple.

Overall, both ILs interact with both conformational states, but with distinct profiles. [Ch][Glu] perturbs the folded state in a largely nonspecific manner (**Fig. 3c vs. 3d**), whereas [Bmim][dca] produces more specific perturbations in the unfolded state (**Fig. 3e vs. 3f**). Conversely, [Ch][Glu]– unfolded and [Bmim][dca]–folded interactions appear more nonspecific.

In the folded state, CSPs with [Ch][Glu] cluster predominantly in the n-Src loop between β2 and β3 (**Fig. 3c**). With [Bmim][dca], CSPs are distributed throughout the sequence, except for residues E2 and I27 (**Fig. 3e**). In the unfolded state, [Ch][Glu]-induced CSPs are not localized (**Fig. 3d**), whereas [Bmim][dca]-induced CSPs concentrate near residues prone to non-native α-helix formation (**Fig. 3f**). Notably, I24–L28 coincide with a local minimum in a hydrophilicity profile (36).

### Stabilization by [Ch][Glu]: Role of electrostatic and hydration

In the folded state in [Ch][Glu] (**Fig. 3c**), CSP magnitudes are generally low and localize to flexible regions, such as the n-Src loop and the RT-loop between β-sheets 1 and 2. These flexible regions are expected to be susceptible to environmental changes and to interact with dissolved ions. Affected residues include acidic (D8, D15, E31, D32, D33, D9) and polar uncharged residues (S18, N29, S34, N35, N57), with only one basic residue (K6) perturbed. This pattern is consistent with mild [Ch]^+^–protein interactions, although interfacial water exclusion to accommodate anion interaction may also contribute.

Comparisons at 0.35 M with NaCl, Na[Glu], and [Ch]Cl (**Fig. S3**) show that the overall [Ch][Glu] CSP pattern resembles [Ch]Cl more than Na[Glu] or NaCl (**Fig. S3a vs. S3c**). Interestingly, residues E2, A3, I27, and L28 display significant CSPs in NaCl, Na[Glu], and [Ch]Cl but not in [Ch][Glu]. These residues are likely to participate in hydrogen bonds (e.g., I27 with E2 and/or A3, and L28 with β3), and their CSPs in simple salts likely reflect electrostatic screening at elevated ionic strength. The absence of these perturbations in [Ch][Glu] suggests distinct IL effects relative to its constituent ions, potentially resulting from protein interactions with contact ion pairs formed by the IL (37).

Two proximal clusters of acidic residues (D14–E16 and E31–D33) are perturbed by [Ch][Glu] (**Fig. 3c**). These clusters participate in inter-protein electrostatic interactions, and their removal destabilizes SH3 due to unopposed electrostatic repulsion (24, 38). Stabilizing these repulsive interactions by interaction with [Ch^+^] could contribute to the net stabilization by [Ch][Glu]. However, as shown previously for Na_2_SO_4_, single/double mutations of acidic residues do not fully stabilize SH3, indicating that shielding alone is insufficient (24). Consistent with **Fig. 2** and **Table 1**, the stabilizing effect of [Ch][Glu] is primarily anion-driven: [Ch][Glu] and Na[Glu] have similar *m*-values, while [Ch]Cl has minimal effect. Thus, the net stabilization must originate predominantly from [Glu]^−^, not from direct [Ch]^+^ binding.

In parallel with partial screening of electrostatic repulsions, the CSPs are also compatible with some ion accumulation at the macromolecule/water interface (15, 18, 39), a “salting-in”–like scenario that would be destabilizing on its own. We propose this is overcompensated by the strongly stabilizing, kosmotropic character of [Glu]^−^, which, due to its strong hydration, reduces surface tension, and makes it energetically unfavorable for nonpolar parts of the protein to become exposed to the bulk solvent, thereby favoring the more compact folded state (40, 41). Similar behavior has been reported for acetate in cholinium ILs (42).

Within the folded state, residues in the n-Src loop (N29, M30, E31, D32, S34, N35) display CSPs consistent with a binding isotherm (**Fig. S3e**), yielding an apparent equilibrium dissociation constant K_D_ ≈ 0.11 M ± 0.05 M. This affinity aligns with the physiological concentration of glutamate in cells (~0.1 M in *E. coli* (43)) and suggests that, *in vivo*, the equilibrium of SH3 may be more shifted toward the folded state than observed *in vitro*. However, the relevance of these interactions in the crowded cellular environment remains to be studied.

Although both [Ch][Glu] and Na_2_SO_4_ stabilize SH3, their CSP signatures differ markedly (**Fig. S4**). In ~1.0 M [Ch][Glu] (~100% folded), perturbations cluster in specific regions (e.g., the n-Src loop), whereas in 0.4 M Na_2_SO_4_ (~100% folded), affected residues are broadly distributed without bias. This points to distinct stabilization mechanisms. The small CSP magnitudes in [Ch][Glu], like those in Na_2_SO_4_, imply that the folded state closely resembles that in water, though subtle structural differences cannot be excluded (*see* following section).

Because 0.35 M [Ch][Glu] does not fully shift the equilibrium to the folded state, we also examined the effects in U_[Ch][Glu]_. CSPs in U_[Ch][Glu]_ (**Fig. 3d**) show that some residues in the unfolded state are perturbed, but – with the exception of W36 and the C-terminal D59 – do not overlap with those in F_[Ch][Glu]_. This indicates that IL–protein interactions depend not only on residue type but also on conformation. The affected residues in U_[Ch][Glu]_ are dispersed and predominantly hydrophobic (9/16), consistent with greater solvent exposure upon unfolding. As in the folded state, the small CSP magnitudes suggest that U_[Ch][Glu]_ is similar to U_water_.

Finally, CSPs induced by Na[Glu] and [Ch]Cl (**Fig. S5**) do not reveal a clear preference for either [Ch]^+^ or [Glu]^−^ alone; the observed pattern appears to result from their combined presence together with ionic-strength effects.

### Destabilization by [Bmim][dca]: Cooperative action of cation and anion

Like [Ch][Glu], [Bmim][dca] interacts with both the folded (F_[Bmim][dca]_) and unfolded (U_[Bmim][dca]_) states of SH3 (**Fig. 3e–g**). In the folded state, ~20/59 residues are perturbed and distributed across the sequence without a strong bias by residue type (five acidic, five polar uncharged, seven hydrophobic, three basic). Most affected residues are surface-exposed, and the fraction of perturbed hydrophobic residues is higher than in [Ch][Glu]. Notably, only three residues (K6, D15, E31) are affected in common between F_[Bmim][dca]_ and F_[Ch][Glu]_, all in unstructured loops. These observations agree with prior reports that [Bmim]^+^ cations bind hydrophobic patches and dehydrate charged environments, thereby destabilizing proteins due to the favorable interactions of the hydrophobic surface of the protein in the unfolded state with the IL (15, 16, 44). The destabilizing behavior is reminiscent of guanidinium ([Gdm]^+^), which also targets hydrophobic surfaces and disrupts stability (45).

CSP analyses with NaCl, [Bmim]Cl, and Na[dca] at 0.15 M (**Fig. S6**) indicate that [Bmim]^+^ accounts for most of the perturbations observed with [Bmim][dca] in the folded state. Remaining CSPs largely reflect electrostatic screening, as seen with NaCl (**Fig. S6b**). Thus, although the strong denaturing power of [Bmim][dca] has been linked to the high H-bond basicity of [dca]^−^ (15, 16, 46), our CSP data suggest that, for the folded state, specific [Bmim]^+^–protein interactions combined with electrostatics dominate the perturbation pattern.

In the unfolded state, the scenario differs markedly (**Fig. 3f**). CSPs are highly localized to Q23– N29, adjacent to a non-native α-helix observed in buffer (26, 35), and encompass hydrophobic residues (I24, L25, I27, L28) that form β2 in the native fold. The lack of significant perturbation of these residues in [Bmim]Cl and NaCl (**Fig. S7b,c**) argues that the effect originates from [dca]^−^ rather than from regional hydrophobicity alone. The CSP pattern implicates [dca]^−^ interactions with K26, promoting backbone dehydration and stabilizing a compact, helical-like conformation in the unfolded ensemble. [Bmim]^+^ also contributes, particularly at Y37 and R38, likely via hydrophobic contacts with the aromatic side chain of Y37. The more hydrophobic cation–anion ion pair provides additional stabilization relative to [dca]^−^ alone, consistent with CSP differences between [Bmim][dca] and Na[dca] (**Fig. S7a,d**).

Overall, the strong destabilization by [Bmim][dca] reflects cooperative cation/anion actions, consistent with prior observations for [Bmim][Br] (47). Comparisons between [Bmim]Cl and Na[dca] in both states (**Fig. S6c,d**; **Fig. S7c,d**) support a division of labor: [Bmim]^+^ primarily perturbs the folded state, whereas [dca]^−^ drives changes in the unfolded state.

To test whether shielding the positive charge at K26 could favor local helicity, we used AGADIR to estimate non-native helical propensities for SH3 mutants K26A, K26G, K26M, and K26Q (26, 48, 49). Sequences were submitted at 298 K and ionic strength 0.15 M (http://agadir.crg.es). As expected for unfolded conformations, helical propensities were low (<7%), yet replacing K26 increased predicted helicity around the mutation site, closely mirroring the CSP-defined hotspot (**Fig. S8**).

Overall, the observed stronger destabilizing properties of this [Bmim][dca] as compared to that of its ionic salts is in line with a cooperative effect of the [Bmim]^+^ cation with the [dca]^−^ anion where the first destabilizes the folded structure via the direct binding to the protein surface while the latter seems to stabilize the unfolded structure via a stabilization of a particular hydrophobic patch, located in a region directly adjacent to the non-native α-helical structure observed in water. This cooperative destabilizing effect is in line with that shown previously for [Bmim][Br] (47). The stabilization of the unfolded state may be further increased by the hydrophobic contacts with the ion-pair, which stabilizes a particular hydrophobic patch.

Consistent with **Fig. S1**, as little as ~0.35 M [Bmim][dca] shifts the equilibrium to nearly 100% unfolded, yet Δδ_comb_ values in the unfolded ensemble continue to depend on IL concentration (**Fig. 3g,h**). A fit to residues I24, L25, K26, I27, and L28 yields an average K_D_ ≈ 0.6 ± 0.1 M, consistent with weak, nonspecific binding.

We also used AGADIR to examine the effect of ionic strength on SH3 helicity (**Fig. S8b**). Increasing salt enhances helical propensity immediately after to the α-helix (the same segment showing strong CSPs) in a concentration-dependent manner, fully consistent with our NMR observations. The region around Y37 retains some helical tendency that is insensitive to K26 mutation (**Fig. S8a**) but is markedly reduced at higher ionic strength (**Fig. S8b**), suggesting greater solvent exposure of this hydrophobic patch and enhanced accessibility to [Bmim]^+^.

In summary, [Bmim][dca] is a stronger denaturant than its component salts because [Bmim]^+^ destabilizes the folded state via direct binding to hydrophobic surfaces while [dca]^−^ stabilizes a specific hydrophobic/helix-prone patch in the unfolded ensemble adjacent to a non-native helix observed in water. Hydrophobic contacts with the ion pair may further enhance stabilization of this patch, reinforcing the cooperative mechanism (cf. [Bmim][Br]; (47).

### Comparison with other denaturants

To compare [Bmim][dca] with guanidinium chloride ([Gdm]Cl), we analyzed U_[Bmim][dca]_ at 1 M and U_[GdmCl]_ at 2 M (**Fig. S9**). In [Bmim][dca], sharp, intense resonances indicate fast exchange within the unfolded ensemble, whereas in [Gdm]Cl, severe line broadening suggests restricted conformational sampling. CSP patterns also diverge: [Bmim][dca] induces localized perturbations (residues 23–30), whereas [Gdm]Cl perturbs more broadly. Prior work showed that [Gdm]Cl destabilizes Q23–L28 by erasing α-helical propensity in favor of random coil (38, 50, 51). In contrast, [Bmim][dca] appears to stabilize residual structure in this region, indicating fundamentally different unfolded ensembles (*vide infra*).

### IL-induced effects on protein structure and conformation

The chemical-shift dependence of the SH3 domain on cosolutes suggests that the structures of F_[Ch][Glu]_ and U_[Bmim][dca]_ deviate, to different extents, from those in water (F_water_ and U_water_). This is especially evident for U_[Bmim][dca]_, where the most strongly perturbed residues lie directly adjacent to the non-native helical segment present in U_water_ (**Fig. 3h–i**).

To quantify structural changes, we assigned backbone and aliphatic chemical shifts (HN, N, Cα, Cβ, CO) for both folded and unfolded states in water (F_water_ and U_water_; **Fig. 4a**) and under conditions that fully stabilize each state – F_[Ch][Glu]_ (**Fig. 4b**) and U_[Bmim][dca]_ (**Fig. 4c**). Assignments are deposited in the BMRB (accession **53285**). We computed secondary-structure propensity (SSP) from Cα/Cβ shifts (52) to estimate fractional α- and β-content. SSP profiles for water (gray), [Ch][Glu] (green), and [Bmim][dca] (purple) were compared for the folded and unfolded ensembles (**Fig. 4d,e**).

**Fig. 4.**
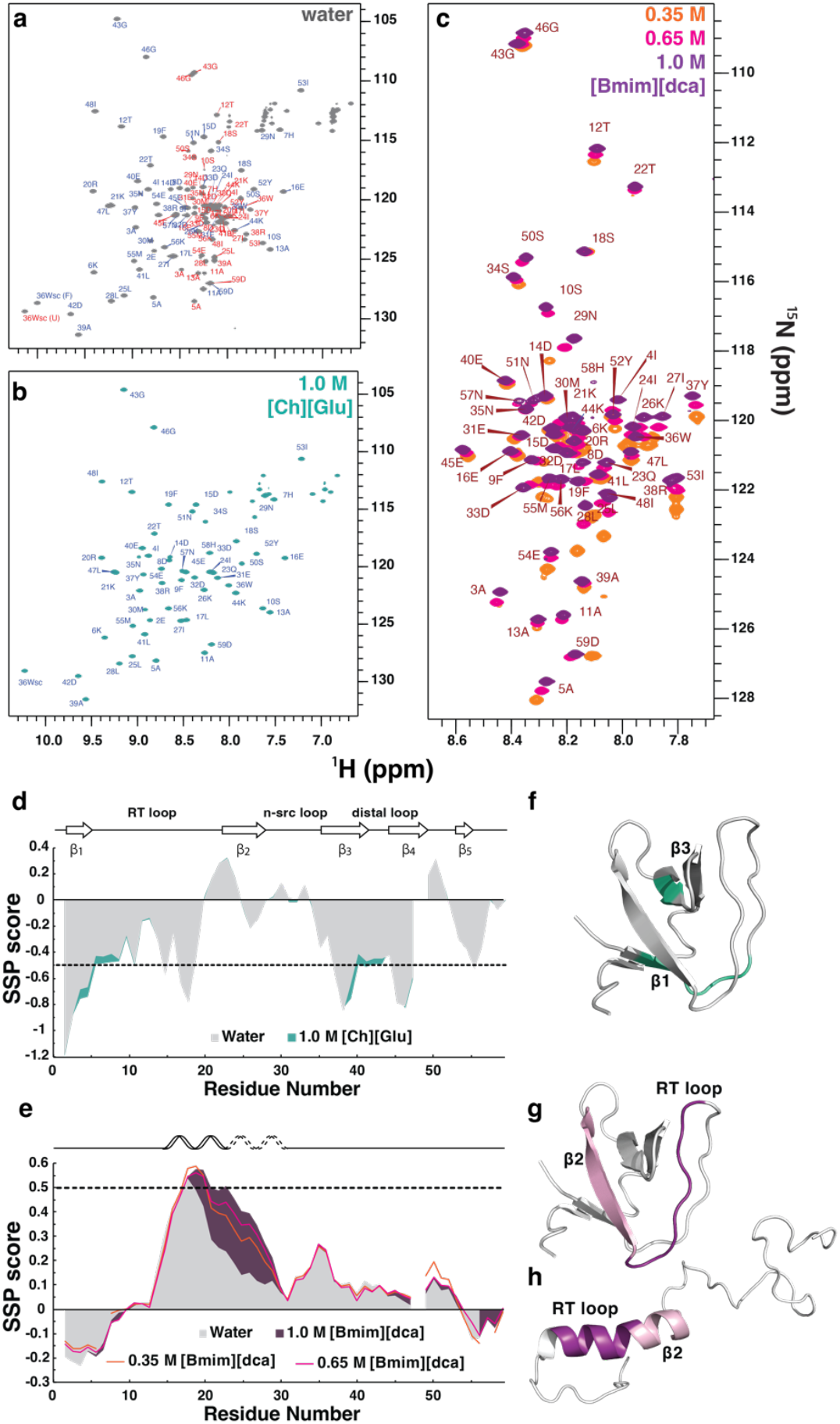
Secondary structure of SH3 in the presence of ILs. **(a)** [^1^H,^15^N]-HSQC of SH3 in water (gray). Blue/red labels denote folded/unfolded resonances. **(b)** Spectrum in 1.0 M [Ch][Glu] (turquoise)—fully folded, stabilized state; assignments in blue. **(c)** Overlay of spectra in 0.35 M (orange), 0.65 M (pink), and 1.0 M [Bmim][dca] (purple) - fully unfolded, stabilized state; assignments shown for 1.0 M. SSP profiles under stabilizing (**d**; [Ch][Glu], turquoise) and destabilizing (**e**; [Bmim][dca], purple) conditions versus water (gray). **(f)** Folded SH3 (PDB: 2A36) highlighting affected regions (turquoise) by [Ch][Glu]. Folded **(g)** and representative unfolded (**h**; PED00022 #1195) structures highlighting β2 (pink) and the RT loop (purple) affected by [Bmim][dca]. SSP was computed from Cα/Cβ shifts; scores of +1/–1 correspond to fully formed α/β, and ±0.5 (dashed lines) to 50% population. Data at 298 K, pH 7.1 in water (gray), 1.0 M [Ch][Glu] (turquoise), and increasing [Bmim][dca] (colors as in **b**). In **(e)**, dashed lines in the secondary structure representation above the plot indicate the potential extension of the α-helix.

For F_[Ch][Glu]_, SSP scores differ only subtly from F_water_ (**Fig. 4d**), consistent with the small-magnitude CSPs (**Fig. 3c,d**). The differences trend toward a slight increase in β-propensity, compatible with a modest stabilization of β-structure without detectable remodeling of the native fold. This mirrors earlier observations for F[Na_2_SO_4_], where NOE patterns and chemical shifts indicated no significant structural deviation from F_water_ (24). Thus, despite small CSPs in [Ch][Glu] and [Na_2_SO_4_], F_water_, F_[Ch][Glu]_, and F[Na_2_SO_4_] appear to be essentially the same.

In contrast, in U_[Bmim][dca]_, residues R20 and N29 show marked CSPs (**Fig. 3f,g**) accompanied by a significant increase in α-helical propensity (**Fig. 4e**). Notably, this α-propensity increases with [Bmim][dca] concentration, even though the protein is already fully unfolded at ~0.35 M. The trend aligns with both CSP data (**Fig. 3e–j**) and AGADIR predictions (**Fig. S8**), indicating that [Bmim][dca] stabilizes – and likely extends – the non-native helix present in water.

A cartoon of a representative unfolded conformer illustrates the helix extension in U_[Bmim][dca]_ (**Fig. 4h**). Relative to the folded state (**Fig. 4g**), the entire β2 segment (pink in the native structure) is converted into an α-helix within the unfolded ensemble. Stabilization of this non-native element is expected to retard productive folding, thereby contributing to the observed shift of the equilibrium toward U in [Bmim][dca].

This hydrophobic segment is highly solvent-exposed and corresponds to a region implicated in chaperone recognition (53, 54). In chaperone-bound or crowded conditions, signals from such regions are typically severely broadened. Aqueous [Bmim][dca] offers a practical workaround: amide resonances remain in fast exchange in [^1^H,^15^N]-HSQC spectra, allowing detection of hydrophobic segments and residual structure that would otherwise be invisible. This resembles denaturant titrations applied to IDPs to mitigate line broadening (55, 56).

In summary, ILs modulate SH3 structure by distinct mechanisms. [Ch][Glu] stabilizes the folded state without detectable conformational remodeling. By contrast, [Bmim][dca] primarily destabilizes by stabilizing a specific non-native α-helical segment within the unfolded ensemble, rather than by directly disrupting the native β-sheet architecture. This unfolded state differs markedly from that observed in [Gdm]Cl, as shown by distinct CSP and SSP signatures (**Fig. S9**). More broadly, these findings underscore the potential of ILs as experimental tools to study defined conformations within unfolded ensembles, including those with possible physiological relevance.

### Thermodynamic interpretation of IL-induced effects

To uncover the thermodynamic bases of [Ch][Glu]- and [Bmim][dca]-induced stability changes, we measured the thermal stability of SH3 (278–313 K) in water and in the presence of 0.35 M [Ch][Glu] or 0.15 M [Bmim][dca] (**Fig. 5a**). The folded (F) and unfolded (U) states of SH3 form a reversible two-state equilibrium (**Eq. 1**). Protein stability is determined by the Gibbs free energy of unfolding, which reflects enthalpy–entropy compensation (**Eq. 3**). A positive value of Δ*G*^0’^_u_ indicates an excess of folded over unfolded:

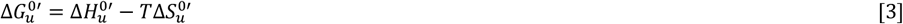

**Fig. 5.**
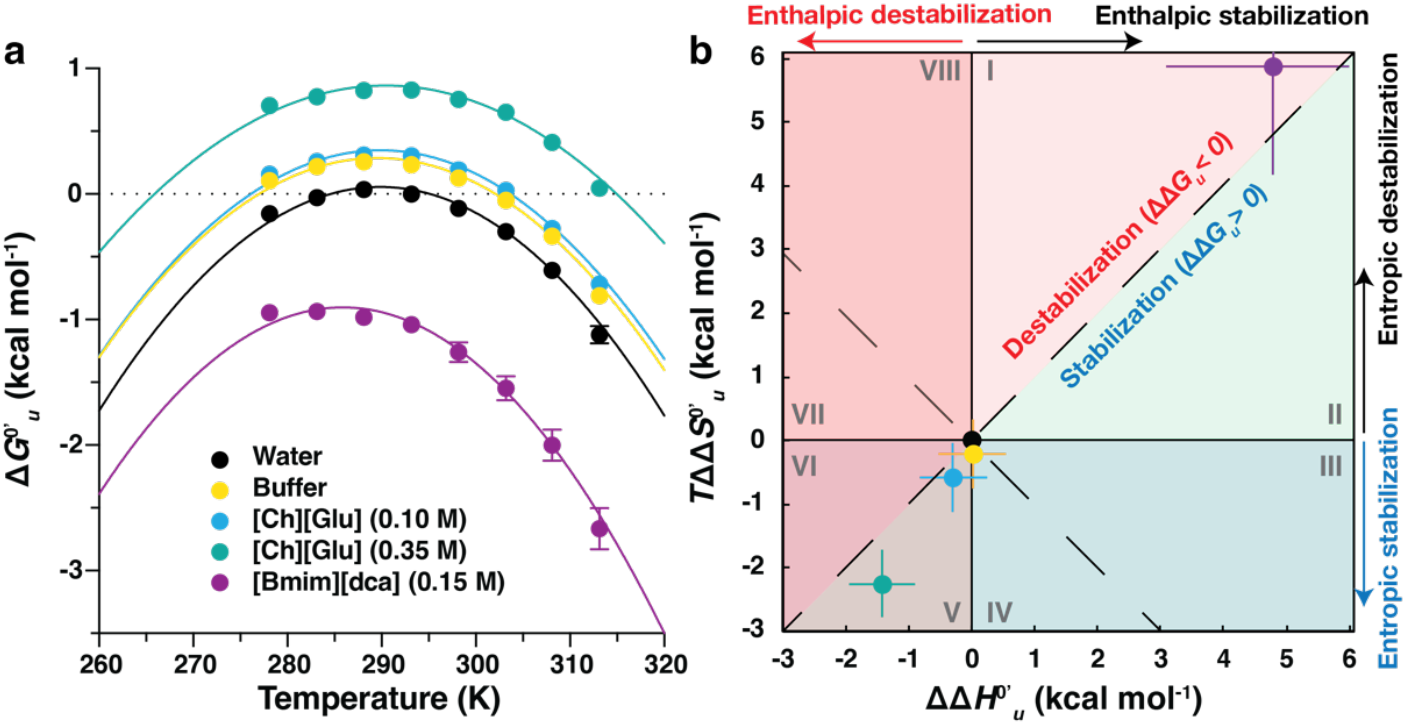
Thermodynamic fingerprint of SH3 in the presence of ILs. (**a**) Stability curves of SH3 in the presence of ILs. Experimental data points for drkN SH3 in water, buffer 0.05 M sodium phosphate pH 7.2, 0.10 and 0.35 M [Ch][Glu], and 0.15 M [Bmim][dca], measured from 278 K to 313 K in 5 K increments. Error bars for almost all conditions are smaller than the labels and represent the propagation of uncertainty in peak volume. The lines correspond to the fitting curves as calculated from **Eq. 4**. Details can be found in the Supporting Information, under the section: SI Materials and Methods. (**b**) Enthalpy-entropy compensation plot. Fields denote combinations of ΔΔ*G*^0’^_u_, ΔΔ*H*^0’^_u_, *T*ΔΔ*S*^0’^_u_ (evaluated at *T*_*m*, water_ ≈ 294 K). Adapted (62, 63). The blue diagonal indicates full enthalpy-entropy compensation and separates the protein destabilizing (ΔΔ*G*^0’^_u_ < 0) from the stabilizing (ΔΔ*G*^0’^_u_ > 0) region. Positive (negative) values of ΔΔ*H*^0’^_u_ imply stabilization (destabilization) by the cosolute. The entropy term acts in the opposite direction: a positive TΔΔ*S*^0’^_u_ contributes to destabilization.

Rather than focusing only on *T*_*m*_ (where Δ*G*^0’^_u_ =0 and *p*_*f*_ = *p*_*u*_ = 0.5), we constructed protein stability curves by measuring Δ*G*^0’^_u_ (*T*) across temperature (57). For globular proteins, ΔG^0’^_u_ (*T*) is typically parabolic with a maximum at *T*_*s*_ (temperature of maximal stability) and zeros at *T*_*m*_ (heat denaturation) and *T*_*c*_ (cold denaturation).

We quantified *p*_*f*_ and *p*_*u*_ from the two W36 indole N^ε1^-H^ε1^ signals (slow exchange [^1^H,^15^N]-HSQC), analogous to established ^19^F-Trp readouts (58–61). From these, we obtained Δ*G*^0’^_u_ (*T*) for SH3 in water, buffer (50 mM sodium phosphate, pH 7.2), 0.10 and 0.35 M [Ch][Glu], and 0.15 M [Bmim][dca] (**Fig. 5a**; **Table S1**). Fitting to the integrated Gibbs–Helmholtz relation:

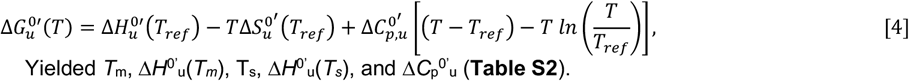

Yielded *T*_m_, Δ*H*^0’^_u_ (*T*_*m*_), T_s_, Δ*H*^0’^_u_ (*T*_*s*_), and Δ*C*_p_ ^0’^_u_ (**Table S2**).

where *T*_*ref*_ is a reference temperature and Δ*C*_*p*,*u*_^*0’*^ is the heat capacity change upon unfolding (Δ*C*_*p*_ ^*0’*^_*u*_ = *C*_*p*_ ^*0’*^_*U*_ – *C*_*p*_ ^*0*^_*F*_^*’*^ ), which we assume is temperature independent over the range studied (278 K to 313 K).

### Baseline

In water, *T*_*m*_ = 294 K, and the buffer slightly stabilizes SH3 relative to water, consistent with previous studies (58, 59).

### [Ch][Glu]

At 0.10 M, stability increases marginally; at 0.35 M, stabilization is pronounced with Δ*G*_0’ u_ (*T*_*m*,*water*_) = 0.83 ± 0.05 kcal mol^-1^. This is accompanied by a significant increase in Δ*H*^0’^_u_ and a decrease in Δ*C*_p_ ^0’^_u_ (ΔΔ*C*_p_ ^0’^_u_ = Δ*C*_p_ ^0’^_u,0.35MChGlu_ ^−^ Δ*C*_p_ ^0’^_u,water_ = −0.3 ± 0.1 kcal mol^-1^ K^-1^), leading to a shallower stability curve and a broader unfolding transition. The lower Δ*C*_p,u_^0’^ implies reduced solvent-accessible surface area in U, consistent with excluded-volume stabilization that restricts configurational entropy of the unfolded ensemble (cf. glucose-induced stabilization; (59)).

### [Bmim][dca]

At 0.15 M, SH3 is destabilized with Δ*G*^0’^_u_ (*T*_*m*,*water*_) = −1.08 ± 0.06 kcal mol^-1^, accompanied by increases in both enthalpic and entropic terms (**Table S2**).

To dissect the thermodynamic mechanisms, we evaluated excess quantities relative to water at *T*_*m*,*water*_ : ΔΔ*G*^0’^_u_, ΔΔ*H*^0’^_u_, and TΔΔ*S*^0’^_u_ (**Eq. 4**; **Table S3**) and mapped them on an enthalpy–entropy compensation diagram (**Fig. 5b**) that partitions eight regimes by the signs/magnitudes of ΔΔ*H*^0’^_u_ and TΔΔ*S*^0’^_u_ (62–64). Stabilizers have: ΔΔ*G*^0’^_u_ > 0 (below diagonal; regions II-V), destabilizers: ΔΔ*G*^0’^_u_ < 0 (above diagonal; regions I, VI-VIII). Positive (negative) ΔΔ*H*_u_^0’^ contributes to stabilization (destabilization), while positive TΔΔ*S*^0’^_u_ drives destabilization.

### [Ch][Glu] is an entropically driven stabilizer

[Ch][Glu] stabilizes SH3 with ΔΔ*G*^0’^_u_ > 0 and *T*ΔΔ*S*^0’^_u_< 0, consistent with excluded-volume effects (65) that preferentially lower the entropy of U relative to F. This behavior parallels hydrophilic cosolutes (inorganic salts such as NaCl/KCl/K_2_SO_4_; sugars like glucose/sucrose (59); osmolytes such as TMAO (66); and high-MW PEGs (61). The concomitant increases in ΔΔ*G*^0’^_u_ and Δ*T*_*m*_ (i.e., Δ*T*_*m*_ = 9 ± 1 at 0.1 M and Δ*T*_*m*_ = 20 ±1 at 0.35 M; **Table S3**) reinforce an entropy-dominated stabilization mechanism.

CSPs (**Fig. 3c,d**) show only interactions with solvent-exposed regions, implying some partitioning to the protein surface. An alternative “preferential exclusion” of [Ch][Glu] ions or ion-pairs scenario (67) would predict preferential hydration and enthalpic stabilization (ΔΔ*H*^0’^_u_ > 0) partially offset by TΔΔ*S*^0’^_u_ > 0; however, our thermodynamic data at 0.35 M [Ch][Glu] indicate TΔΔ*S*^0’^_u_ < 0, favoring the excluded-volume interpretation (**Tables S2–S3**; **Fig. 5**).

Related systems (e.g., KH_2_PO_4_ and [Ch][H_2_PO_4_]) display a concentration-dependent switch from enthalpic to entropic stabilization in the 0.25–0.5 M regime, coincident with maxima in ΔΔ*H*_u_^0’^ and *T*ΔΔ*S*^0’^_u_ (63), often interpreted as a transition from nonspecific electrostatics to ion-specific interactions (62, 68). While our [Ch][Glu] dataset does not span that transition, such behavior may underlie non-monotonic *T*_*m*_ trends reported at high salt and for cholinium ILs solutions (69).

### [Bmim][dca] is an entropically driven destabilizer with partial enthalpic compensation

For [Bmim][dca], ΔΔ*G*^0’^_u_ < 0 and both ΔΔ*H*^0’^_u_ and *T*ΔΔ*S*^0’^_u_ > 0. Thus, unfolding becomes more endothermic (enthalpic stabilization of F), but the larger positive entropic term drives net destabilization. The decrease in the compensation temperature (Δ*T*_*s*_ = ^−^4 ± 1 K; **Table S3**) supports an entropy-dominated mechanism. Unlike classical denaturants (urea, [Gdm]Cl) that destabilize via strong enthalpic interactions with the backbone (ΔΔ*H*^0’^_u_ < 0;(70)), [Bmim][dca] acts differently: it raises both ΔΔ*H*^0’^_u_ and *T*ΔΔ*S*^0’^_u_ and lowers stability because the entropic term dominates.

A plausible molecular picture is that exposure of hydrophobic regions upon unfolding incurs a smaller entropy penalty in [Bmim][dca] (bulk water already more structured/entropy-reduced), yielding *T*ΔΔ*S*^0’^_u_ > 0. Simultaneously, ΔΔ*H*^0’^_u_ > 0 implies unfolding is more enthalpically costly – consistent with less favorable solvation of exposed backbone in the IL medium – providing only partial compensation (71). The butyl chain of [Bmim]^+^ and its synergy with [dca]^−^ likely amplify this effect. Importantly, our structural analysis (*see* preceding section) indicates that [Bmim][dca] destabilizes indirectly by stabilizing a non-native α-helical segment in U rather than by directly disrupting the native β-sheet, expanding the mechanistic repertoire of IL-driven denaturation.

### ILs reshape the folding landscape: kinetics and transition-state energetics

To probe how [Ch][Glu] and [Bmim][dca] alter the SH3 folding landscape, we measured interconversion rates between the folded and unfolded ensembles using longitudinal nitrogen magnetization exchange NMR (ZZ-exchange, ZZex) (22, 72). The SH3 domain undergoes slow two-state exchange with *k*_*ex*_ ≈ 2.2 s^−1^ under near-native conditions (50 mM sodium phosphate, pH 6.0, 293 K) (22). In ZZex spectra, auto-peaks (*ff, uu*) and exchange cross-peaks (*fu, uf*) report on magnetization transfer during the mixing time; their time dependence follows the longitudinal Bloch–McConnell equations, yielding *k*_*ex*_, folding (*k*_*f*_), and unfolding (*k*_*u*_) rates, as well as longitudinal ^15^N relaxation rates (R_1f_ and R_1u_) (22, 73).

We recorded ZZex datasets at 293 K for SH3 in water, 0.35 M [Ch][Glu], and 0.15 M [Bmim][dca], conditions under which *p*_*f*_ ≈ *p*_*u*_ ≈ 0.5 in water, *p*_*f*_ ≈ 0.8 in [Ch][Glu], and *p*_*f*_ ≈ 0.2 in [Bmim][dca]. Example intensity time-courses for T22 are shown in **Fig. 6a–e**. Representative spectra are shown in **Figs. S10** and **S11**. Residue-specific changes in δ*ω*_UF_ between water and ILs (**Fig. S12**) mirror the CSP hotspots (**Fig. 3**), notably I27 and R38 in [Ch][Glu] and I24–L28 in [Bmim][dca]. Fit parameters (*p*_*f*_, *p*_*u*_, R_1f_, R_1u_, *k*_*u*_ and *k*_*f*_) appear in **Table S4**.

**Fig. 6.**
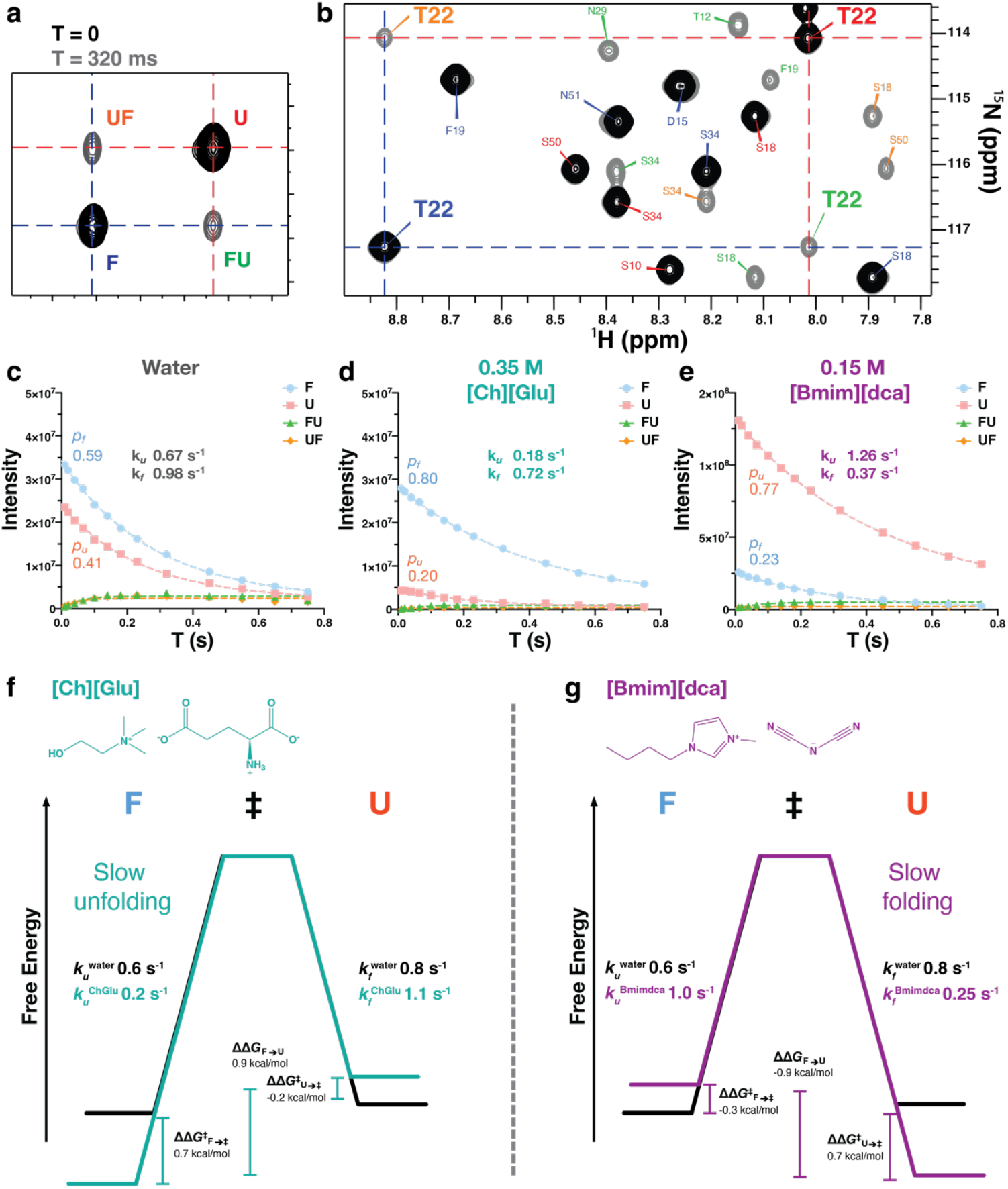
ZZ-exchange experiment and derived models of SH3 free energy landscape. **(a)** Diagram illustrating ZZex typical spectra for a ^1^H/^15^N spin pair undergoing slow two-site exchange within a mixing period T. Blue/red: F/U auto-peaks; green/orange: FU/UF exchange peaks produced during time *t*. **(b)** Region of the ^15^N/^1^H ZZex-HSQC showing T22 for SH3 in water, 293 K, 600.13 MHz; auto-peaks (black), cross-peaks (gray), T = 320 ms. **(c–e)** Intensity time-courses for T22 in water, 0.35 M [Ch][Glu], and 0.15 M [Bmim][dca]; dashed lines are best fits for magnetization initially in F (ff→fu) or U (uu→uf). **(f)** Free-energy diagram for [Ch][Glu] showing increased F→TS‡ barrier (slower *k*_*u*_) and mildly increased *k*_*f*_. **(g)** Diagram for [Bmim][dca] showing decreased U→TS‡ barrier (slower *k*_*f*_) and increased *k*_*u*_. Rates labeled *k*_*f*_ and *k*_*u*_ are viscosity-corrected averages (Table 2; see mapping above). ΔΔ*G*^0’^_u_, ΔΔ*G*^*0’‡*^ _*F U*→*TS‡*_, are mean values with propagated uncertainties.

### Viscosity correction

Because diffusion-controlled rates scale as 1/η, we corrected measured rates by the relative viscosity *η*_rel_ to obtain intrinsic values (*see* **SI Material and Methods**). To avoid ambiguity, we use: *k*_*f*_ ≡ *k*_U→F_, *k*_*u*_ ≡ *k*_F→U_.

**Table 2.** Average parameters for the SH3 interconversion extracted from ZZex.

| | $p_f$ | $p_u$ | $R_{1f}$<br>(s<sup>−1</sup>) | $R_{1u}$<br>(s<sup>−1</sup>) | $k_u$ (s<sup>−1</sup>) | $k_f$ (s<sup>−1</sup>) | $k_{ex}$ (s<sup>−1</sup>) |
| --- | --- | --- | --- | --- | --- | --- | --- |
| **Water** | 0.54 ± 0.08 | 0.46 ± 0.08 | 2.5 ± 0.2 | 2.3 ± 0.3 | **0.6 ± 0.1** | **0.8 ± 0.2** | **1.4 ± 0.2** |
| **0.35 M**<br>**[Ch]**<br>**[Glu]** | 0.83 ± 0.05 | 0.17 ± 0.05 | 2.1 ± 0.1 | 2.2 ± 0.3 | 0.16 ± 0.07 | 0.8 ± 0.3 | 1.0 ± 0.3 |
|  |  |  |  |  | **0.2 ± 0.1<sup>a</sup>** | **1.1 ± 0.4<sup>a</sup>** | **1.3 ± 0.3<sup>a</sup>** |
| **0.15 M**<br>**[Bmim]**<br>**[dca]** | 0.20 ± 0.07 | 0.80 ± 0.06 | 2.6 ± 0.5 | 2.3 ± 0.2 | 1.0 ± 0.3 | 0.24 ± 0.09 | 1.3 ± 0.3 |
|  |  |  |  |  | **1.0 ± 0.3<sup>a</sup>** | **0.25 ± 0.09<sup>a</sup>** | **1.3 ± 0.3<sup>a</sup>** |

In **Table 2**, the columns corresponding to *k*_*u*_ and *k*_*f*_, after viscosity correction (*η*_rel_ =1.39 for 0.35 M [Ch][Glu]; *η*_rel_ =1.02 for 0.15 M [Bmim][dca]), satisfy the expected steady-state relation *p*_*f*_ ≈ *k*_*f*_ /(*k*_*f*_+*k*_*u*_)*p*_*f*_ within experimental dispersion.

Interconversion rates were analyzed for 15 residues under each of the three experimental conditions. The residues are well distributed across the sequence (S10, T12, A13, E16, S18, T22, L25, L28, N29, A39, D42, G46, I53, K56, and W36sc; **Fig. S13**) and only residues with well-resolved peaks and reliable rates across all three conditions were included in the analysis. Viscosity-corrected rates are provided in **Table S5**; and the mean values are summarized in **Table 2**.

The average from 15 residue values extracted from fitting for SH3 in water is 0.35 M [Ch][Glu], and 0.15 M [Bmim][dca] at 293.2 K, 600.13 MHz. Uncertainties represent the standard deviation from the average. ^a^ Viscosity-corrected rates, the relative viscosity used for 0.35 M [Ch][Glu] and 0.15 M [Bmim][dca] was 1.39, and 1.02, respectively (see **SI Materials and Methods**).

### Opposing kinetic signatures of [Ch][Glu] and [Bmim][dca]

[Ch][Glu] stabilizes the folded ensemble by increasing *k*_*f*_ and decreasing *k*_*u*_ (viscosity-corrected averages: *k*_*f*_ = 1.1 ± 0.4 s^−1^ *vs* 0.8 ± 0.2 s^−1^ in water; *k*_*u*_ = 0.20 ± 0.10 s^−1^ *vs* 0.60 ± 0.10 s^−1^ in water). Thus, folding accelerates by ~1.4× while unfolding slows by ~3×, consistent with the thermodynamic stabilization and the larger *p*_*f*_ (**Table 2**). Conversely, [Bmim][dca] stabilizes the unfolded ensemble by decreasing *k*_*f*_ and increasing *k*_*u*_ (viscosity-corrected *k*_*f*_ = 0.25 ± 0.09 s^−1^, *k*_*u*_ = 1.0 ± 0.3 s^−1^). Folding slows by ~3.2× and unfolding speeds up by ~1.7× relative to water, matching the shift to *p*_*f*_ ≈ 0.2. Together with the preceding sections, these kinetic effects support a mechanism in which [Ch][Glu] favors compaction of U, whereas [Bmim][dca] stabilizes a specific non-native helical segment within U, impeding productive folding.

### IL-Induced effects on transition state (TS^‡^) energetics

Using ZZex-derived rates (*k*_*f*_ and *k*_*u*_), we estimated activation free energies for folding (Δ*G*^*0’‡*^_*U*→*TS‡*_) and unfolding (Δ*G*^*0’‡*^_*F*→*TS‡*_^*’*^) via transition-state theory, assuming that interconversion between F and U states is limited by the formation of a transition state (TS^‡^), representing the highest free energy ensemble along the folding pathway, by the Eyring-Polanyi equation (74, 75)

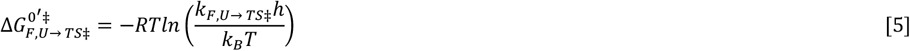

For clarity, *k*_*u*_ = *k*_*F*→ *TS‡*_, *k*_*f*_ = *k*_*U*→ *TS‡*_, and Δ*G*^0’^ = Δ*G*^0’^_u_, with equilibrium stability given by:

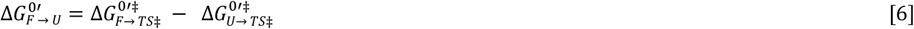

Per-residue activation parameters are listed in **Table S6**; averages are summarized in **Table S7**.

Viscosity effects were also taken into account, since (76) shows that drkN SH3 folding is linearly dependent on solvent viscosity. Viscosity corrections (detailed calculations are described in the **SI Materials and Methods**) lower both Δ*G*^*0’‡*^_*F*→ *TS‡*_ and Δ*G*^*0’‡*^_*U*→ *TS‡*_ modestly in 0.35 M [Ch][Glu] and are negligible for 0.15 M [Bmim][dca]; the qualitative conclusions are unchanged.

Relative to water, [Ch][Glu] increases the unfolding barrier (Δ*G*^*0’‡*^_*F*→ *TS‡*_) with little change in the folding barrier (Δ*G*^*0’‡*^_*U*→ *TS‡*_), yielding a ~3× decrease in *k*_*u*_ and ~1.4× increase in *k*_*f*_ (**Table S7**). This pattern is characteristic of excluded-volume–dominated stabilization: compaction of U and favorable release of interfacial water upon folding raise the F→TS^‡^ barrier while leaving the TS^‡^ itself largely unaltered.

In contrast, [Bmim][dca] lowers Δ*G*^*0’‡*^_*U*→ *TS‡*_, and raises Δ*G*^*0’‡*^_*F*→ *TS‡*_, producing a ~3.2× decrease in *k*_*f*_ and ~1.6–1.7× increase in *k*_*u*_ (**Table S7**). The kinetic signature aligns with the section: *IL-induced effects on protein structure and conformation*: [Bmim][dca] stabilizes a helix-prone, hydrophobic segment in U (I24–L28), which lowers the U→TS^‡^ barrier (U becomes “closer” to TS^‡^) and raises the F→TS^‡^ barrier (F becomes “farther”), causing net destabilization. Thermodynamically, this corresponds to positive ΔΔ*H*^0’^_u_ and positive *T*ΔΔ*S*^0’^_u_ (**Fig. 5b**), unlike urea/[Gdm]Cl, which destabilizes via strong enthalpic binding to the backbone (59, 77).

Although [Bmim][dca] increases helical propensity in U, it does not create a distinct, long-lived helical state; NH resonances remain in fast exchange (**Fig. 4c**). The ensemble remains dynamic, with rapid interconversion among conformers whose average free energy is lowered – similar to SH3 mutants with elevated local helicity that slow folding (26, 78).

Assuming TS^‡^ energy is unchanged, the kinetic/thermodynamic effects can be visualized as in **Fig. 6f,g**: [Ch][Glu] raises the F→TS^‡^ barrier (slower unfolding; stabilized F), while [Bmim][dca] lowers the U→TS^‡^ barrier (slower folding; stabilized U).

### Transition state sensitivity to IL-induced perturbations

Although our results suggest that the most considerable differences in SH3 folding and unfolding rates result from changes in the F or U state energies, we must also consider potential changes in the energy level of the transition state (TS^‡^) (**Fig. S14**). To gauge how much TS^‡^ shifts relative to F and U, we used the proportionality constant α (76, 79):

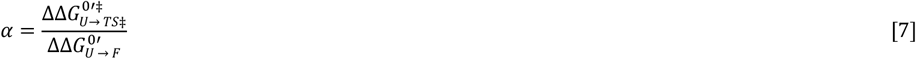

Here, α measures how strongly the U→TS^‡^ barrier responds compared with the overall stability change; larger α often implies a more solvent-exposed, U-like TS^‡^.

Using viscosity-corrected averages from the 15 residues (**Table 2, Table S5-S7**), we find: *α*_[Ch][Glu]_= 0.2 ± 0.6 and *α*_[Bmim][dca]_= 0.7 ± 0.3.

*α*_[Bmim][dca]_ is comparable to values for ~1 M urea or ~1.4 M glycerol (*α*_[Glycerol]_ ≈ *α*_[urea]_ ≈ 0.75 ± 0.1) (76), indicating a partially hydrated TS^‡^ in [Bmim][dca] and suggesting that the non-native helix is partly formed at TS^‡^ (a more “native-like” TS^‡^ along this local coordinate). In contrast, the low α_[Ch][Glu]_ implies that TS^‡^ is not significantly destabilized and may be slightly stabilized, consistent with an excluded-volume mechanism in which the primary energetic shifts occur for F and U rather than TS^‡^. These interpretations assume that rate changes arise from TS^‡^ shifts relative to U (**Fig. S14a,b**).

Because native structural propensities can have multiple origins, we also consider alternative schemes (e.g., **Fig. S14b,c**, where F is unaffected). The convergence of kinetic, thermodynamic, and structural readouts argues that a full accounting of the unfolded ensemble is essential for understanding IL-modulated protein stability.

## Conclusions

In this study, we combined different site-resolved NMR information - chemical shift perturbations (CSPs), secondary-structure propensity (SSP) from Cα/Cβ shifts, ZZ-exchange (ZZex) kinetics, and temperature-dependent stability curves from relative F/U populations quantified through [^1^H,^15^N]-HSQC - to dissect how two ionic liquids reshape the SH3 folding landscape.

Our results demonstrate that the ILs cholinium glutamate ([Ch][Glu]) and imidazolium dicyanamide ([Bmim][dca]) generate orthogonal responses, modulating protein stability through fundamentally different molecular mechanisms that act on distinct regions of the folding landscape. The biocompatible [Ch][Glu] stabilizes SH3 through an entropy-dominated, anion-driven mechanism in which preferential exclusion from the protein surface raises the barrier to unfolding, akin to the action of osmolytes. In contrast, [Bmim][dca] destabilizes SH3 through an entropy-dominated mechanism with partial enthalpic compensation, favoring the unfolded ensemble by stabilizing a compact, non-native α-helical conformation and thereby lowering the barrier to folding. This reveals that unfolded-state stabilization - not just destabilization of the native fold - is a critical determinant of protein conformational equilibria.

These observations challenge the prevailing assumption that cosolutes primarily affect folded states, highlighting the need to consider unfolded ensembles when evaluating protein stability in complex solvents. More broadly, they reveal that ILs are not passive modifiers of ionic strength, but active reshapers of protein energy landscapes, with effects that extend beyond the sum of their individual ions.

By identifying distinct pathways through which ILs stabilize or destabilize proteins, our study provides conceptual and mechanistic foundations for the rational design of ILs tailored to control protein behavior. Such principles hold relevance not only for biocatalysis and therapeutic protein formulation but also for understanding how non-native environments shape protein misfolding, aggregation, and disordered-state function.

## Materials and Methods

Detailed protocols for the synthesis and characterization of choline glutamate ([Ch][Glu]) IL (**Fig. S15**), the expression and purification of ^15^N- and ^15^N/^13^C-labeled drkN SH3, full experimental details, including NMR sample preparation, NMR data acquisition, processing, analysis, and extrapolations, are provided in the **SI Materials and Methods**.

Briefly, [Bmim][dca] and [Bmim][Cl] ILs (≥98% purity, IoLiTec, Germany) were dried under vacuum at 60 °C for 24 hours before use, and [Ch][Glu] was synthesized following a modified protocol from (80) with Amberlyst A-26 ion exchange resin (Thermo Fisher Scientific). The physicochemical properties of [Ch][Glu] were characterized and found to be consistent with previously reported values (81).

The drkN SH3 domain was overexpressed and purified as described by (24).

NMR data were acquired on a Bruker Avance III 600 MHz spectrometer equipped with a TCI cryoprobe. Preliminary drkN SH3 assignments were obtained from the BMRB 25501 (53).

Viscosity measurements of aqueous [Ch][Glu] solutions (**Table S8**) were performed using an SVM 3000 Stabinger viscometer (Anton Paar).

### Data, Materials, and Software Availability

NMR data have been deposited in the Biological Magnetic Resonance Data Bank (BMRB entry: 53285).

## ACKNOWLEDGMENTS

We kindly acknowledge Prof. Dr. Julie Forman-Kay and Dr. João Teixeira (Hospital for Sick Children, Toronto, Canada) for providing us with the SH3 unfolded ensemble of structures (PED8AAC/PED00022) and Prof. Dr. Lewis Kay (University of Toronto, Toronto, Canada) for providing the pET-11d plasmid containing the gene encoding for WT drkN SH3 domain. We thank Dr. Tiago Cordeiro (ITQB-NOVA, Oeiras, Portugal) and Prof. Manuel Prieto (IST, Lisbon, Portugal) for helpful discussions.

This work was supported by Fundação para a Ciência e a Tecnologia (FCT-MCTES), Portugal, through grants PD/BD/128202/2016 (to M.S.S.), PD/BD/148028/2019 (to S.S.F.), 2021.05564.BD (to P.O.), and contracts PINFRA/22161/2016-1 and 2020.00043.CEECIND (to A.V.); the Applied Molecular Biosciences Unit (UCIBIO), which is financed by national funds from FCT-MCTES (UID/ Multi/04378/2019). The NMR spectrometers are part of the National NMR Network (PTNMR), supported by Fundação para a Ciência e a Tecnologia (ROTEIRO/0031/2013 - PINFRA/22161/2016), co-financed by FEDER through COMPETE 2020, POCI, and PORL and FCT through PIDDAC. A.M.F. acknowledges support from the University of Coimbra through contract IT074-22-12388 and Fundação para a Ciência e a Tecnologia (FCT-Portugal) for the funding of the project 2024.07359.IACDC (https://doi.org/10.54499/2024.07359.IACDC) within the scope of PRR financing, the investment RE-C05-i08-Ciência Mais Digital, and Coimbra Chemistry Centre - Institute of Molecular Sciences (CQC-IMS), which is supported by the Fundação para a Ciência e a Tecnologia (FCT), Portuguese Agency for Scientific Research. CQC is funded by FCT through projects UID/PRR/00313/2025 (https://doi.org/10.54499/UID/PRR/00313/2025) and UID/00313/2025 (https://doi.org/10.54499/UID/00313/2025) and IMS through special complementary funds provided by FCT (project LA/P/0056/2020 https://doi.org/10.54499/LA/P/0056/2020).

