## Supporting Information for "NMR reveals specific remodelling of protein folding landscapes in ionic liquids"

Author affiliations: [a]UCIBIO, Chemistry Department, NOVA School of Sciences and Technology, Universidade NOVA de Lisboa, 2829-516, Caparica, Portugal; [b]Associate Laboratory i4HB - Institute for Health and Bioeconomy, NOVA School of Science and Technology, Universidade NOVA de Lisboa, 2819-516 Caparica, Portugal; [c]Coimbra Chemistry Centre, Institute of Molecular Sciences (CQC-IMS), Department of Chemistry, Faculty of Science and Technology, University of Coimbra, Coimbra, 3004-531 Portugal;

[1]Present address: Department of Pharmaceutical Sciences, University of Vienna, Vienna A-1090, Austria; [2]Present address: Department of Molecular Life Sciences, University of Zurich, Zurich, Switzerland.

**This PDF file includes:**

Supporting text
Figures S1 to S15
Tables S1 to S8
SI References

### SI Materials and Methods

Description of chemicals and materials, the synthesis and characterization of [Ch][Glu] IL, and a detailed protocol for expression and purification of $^{15}N$ or $^{15}N$/ $^{13}C$ isotopically labelled drkN SH3 can be found in this section. NMR sample preparation, data acquisition, processing, and analysis procedures, such as quantification of the F/U population, NMR temperature scan analysis, and ZZex data, including thermodynamic and kinetic examination, are described in the sections. Further details on viscosity measurements and extrapolations are also included.

**Chemicals and Materials.** ʟ-glutamic acid (> 98.5 % of purity) was purchased from PanReac. 1-butyl-3-methylimidazolium dicyanamide ([Bmim][dca]) and 1-butyl-3-methylimidazolium chloride ([Bmim][Cl]) ILs were sourced from IoLiTec (Denzlingen, Germany). The ILs were at least 98% pure and were dried for 24 h under vacuum at 60 ºC before solution preparation. Isotopically enriched chemical compounds, [$^{15}N$]-$NH_4Cl$, [$^{13}C$]-glucose and deuterium oxide ($D_2O$) were purchased from Cambridge Isotope Laboratories. Sodium-ʟ-glutamate, ion exchange resin Amberlyst A-26 (OH) and SnakeSkin 3.5K MWCO dialysis tubing were purchased from Thermo Fisher Scientific. Unless otherwise described, all other chemicals were purchased from Sigma-Aldrich. Pure *MilliQ* water with a resistance of 18.4 M*Ω* $cm^{-1}$ was used in all experiments. pH values are direct meter readings uncorrected for any isotope effect and were measured with Docu-pH meter (Sartorius) calibrated with standard solutions.

**Synthesis and characterization of [Ch][Glu] IL.** For the choline glutamate ([Ch][Glu]) IL synthesis, we used, with slight modifications, the method reported by (1). Based on a potentiometric titration *via* neutralization reaction, [Ch][Glu] was synthesized as follows (**Fig. S15**). Choline hydroxide ([Ch][OH]) aqueous solution was prepared from choline chloride ([Ch][Cl]) (4.75 g, 34 mmoles) in water that was passed slowly through a column packed with ~ 43 mL of anion exchange (Amberlyst A-26) resin and then washed. The absence of chloride content was tested with a silver nitrate qualitative test. Freshly [Ch][OH] aqueous solution (175 mL, 31 mmoles) was added dropwise in an equimolar ratio to ʟ-glutamic acid (4.5 g, 31 mmoles) in water. The reaction was monitored by reading the solution pH until it reached the equivalence point (pH=7.00) and the stoichiometry 1:1 [Ch]/[Glu] ratio was confirmed by characterization of the solution by $^{1}H$ NMR using the integral ratio. The mixture was stirred at about 25 °C for 24 h in the dark and subsequently the solvent was evaporated under reduced pressure. The product was dried *in vacuo* for 24 h at 60 °C to yield [Ch][Glu] (8.5 g, >99%) as a slightly yellow oily or glassy compound at room temperature.

The structure of the resulting [Ch][Glu] IL was confirmed by $^{1}H$ and $^{13}C$ NMR spectroscopy (see below the chemical shifts). The spectra were recorded at 25 ºC on a 400 MHz Bruker AVANCE II+ instrument operating at 400.15 MHz for protons and 100.6 MHz for $^{13}C$, equipped with a 5 mm high-resolution BBO probe with pulsed gradient units. Solution was prepared by dissolving [Ch][Glu] in 99.9% $D_2O$ to a concentration of 0.5 M. The glass transition ($T_g$) and decomposition temperature ($T_d$) of pure [Ch][Glu] IL were determined to -21.3 ºC and 221.3 ºC, respectively. The measurements were conducted using a differential thermal analyzer DSC 131 (Setaram) and a thermogravimetric analyzer Labsys EVO (Setaram). These physicochemical properties are consistent with previously reported values (2).

$^{1}H$ NMR (400.15 MHz, $D_2O$, 25 ºC) $\delta_H$ (ppm): 1.84 – 2.02 (m, 2H, $CH_2$, Glu), 2.20 (apparent q, 2H, $CH_2$, Glu), 3.05 (s, 9H, $CH_3$, $CH_3$, $CH_3$, Ch), 3.37 (apparent t, 2H, $CH_2$, Ch), 3.60 (q, J = 4.87, 7.12 Hz, 1H, CH, Glu), 3.88 – 3.93 (m, 2H, $CH_2$, Glu).

$^{13}C$ NMR (100.6 MHz, $D_2O$, 25 ºC) $\delta_C$ (ppm): 27.34 ($CH_2$, Glu), 33.6 ($CH_2$, Glu), 53.90 ($CH_3$, $CH_3$, CH3, Ch), 54.8 (CH, Glu), 55.61 ($CH_2$, Ch), 67.45 ($CH_2$, Ch), 175.1 (CO, Glu), 181.36 (CO, Glu).

**Protein expression and purification.** $^{15}N$ or $^{15}N$/ $^{13}C$ isotopically enriched, N-terminal Src homology 3 (SH3) domain of *Drosophila* signal transduction protein drk (drkN SH3) containing 59 residues, was overexpressed and purified as previously described (3). *Escherichia coli* BL21 (DE3) competent cells (NZYTech) were transformed with the pET-11d plasmid vector containing the gene encoding drkN SH3 protein by heat shock. Cells were grown with shaking at 37 ºC and 180 rpm in M9 minimal medium supplemented with 100 µM $FeSO_4$, 100 µM $CaCl_2$, 2 mM MgSO4, 10 mg/L Thiamine-HCl, 0.5% MEM Vitamins, 1.5 g/L of $^{15}NH_4Cl$ and 4 g/L of glucose or $^{13}C$-glucose, and 100 mg/L ampicillin. Protein expression was induced by adding 1 mM of isopropyl-β-D-thiogalactopyranoside (IPTG, NZYTech) at $OD_{600}$ (optical density at 600 nm) = 0.7. After 2h induction in the same conditions, cells were harvested [4,425 x *g*, 30 min, 4ºC in a JA-10 rotor (Avanti J-26S XPI, Beckman Coulter)] and frozen at -20 ºC overnight. Pellet was resuspended in 30 mL of lysis buffer per liter of culture [50 mM

Tris-HCl (pH 7.5), 2 mM EDTA, 7 mM β-mercaptoethanol and protease inhibitor cocktail (cOmplete ULTRA tablets, Roche)], and then lysed by sonication on ice [10 min at 80% amplitude using 1 min on/off pulse program (UP100H ultrasonic processor, Hielscher)] and centrifuged [30,000 x *g*, 40 min, 4ºC in a JA-25.50 rotor (Avanti J-26S XPI, Beckman Coulter)]. Supernatant was dialyzed (SnakeSkin 3.5K MWCO) overnight at 4ºC against Buffer A [50 mM Tris-HCl (pH 7.5), 2 mM EDTA, and 7 mM β-mercaptoethanol]. The dialyzed sample was loaded to a HiTrap Q HP anion-exchange column (GE Healthcare) on an ÄKTA start system (GE Healthcare), and buffer B [50 mM Tris-HCl (pH 7.5), 1 M NaCl, 2 mM EDTA and 7 mM β-mercaptoethanol] was used to produce a gradient 0 - 500 mM NaCl where drkN SH3 elutes at around 150 mM NaCl since the binding with the column is weak.

Fractions containing the protein of interest were pooled and concentrated with a Vivaspin Turbo 15 3 K MWCO centrifugal concentrator (Sartorius). In a final step, a size exclusion chromatography step [Superdex 75 10/300 GL column (GE Healthcare) in Shimadzu prominence machine, at 4ºC] was applied to further purify and adjust drkN SH3 to a different buffer [50 mM phosphate buffer (pH 7.2) and 150 mM NaCl]. Purity was analyzed in each step by SDS-PAGE (BioRad). Pure samples were extensively desalted by dialysis in *MilliQ* water, flash frozen, lyophilized (Edwards Modulyo Freeze Dyer), and stored at -20 ºC until usage. The final yield of purified protein was 7 mg of drkN SH3 per liter of M9 minimal medium. Concentrations were assessed spectrophotometrically by absorption measurements at 280 nm ($\varepsilon$= 8,480 $M^{-1}$ $cm^{-1}$) using NanoDrop ND-1000 UV-Vis (Thermo Fisher Scientific).

**NMR spectroscopy.** Sample preparation, data acquisition and processing. All samples for NMR spectroscopy were prepared in *MilliQ* water with 10% (v/v) $D_2O$, 0.1% (v/v) $NaN_3$, and 50 μM sodium-2,2-dimethyl-2-silapentane-5-sulfonate-$d_6$ (DSS, Eurisotop). The pH was adjusted to 7.1 ± 0.1 with negligible microliter addition of HCl or NaOH solutions. Except where stated, spectra were recorded at 298.2 K on a 600 MHz Bruker Avance III spectrometer operating at a proton Larmor frequency of 600.13 MHz equipped with a 5-mm TCI cryoprobe. Spectrometer temperature was calibrated using a pure methanol-*d₄* standard Bruker sample (4). Further specific details of data acquisition (pulse programs, spectral windows, etc) and processing for each experiment are given in the following subsections. Proton chemical shifts were referenced against internal DSS, while nitrogen and carbon chemical shifts were referenced indirectly to DSS using the absolute frequency ratio (5). Processing and visualization of NMR spectra were performed with NMRPipe (version 10.1) (6) and CcpNmr Analysis 2.5 (7). Preliminary drkN SH3 assignments were taken from the BMRB databank under the accession code 25501 (8).

NMR Chemical Shift Titrations. The standard Bruker $^1H$-$^{15}N$ HSQC (hsqcetfpf3gpsi2) pulse sequence employs a sensitivity-enhanced pulse field gradient (9). The spectra were acquired with 2048 ($^1H$) and 128 ($^{15}N$) complex points for a spectral width (SW) of 12 ppm ($^1H$) and 32 ppm ($^{15}N$), with 8 scans. Two-dimensional $^1H$-$^{15}N$ HSQC experiments were subsequently collected for different ionic liquids ([Ch][Glu], [Bmim][dca], [Bmim][Cl]) and salts ([Na[Glu]], Na[dca], [Ch]Cl, NaCl) titrations with 350 μM $^{15}N$-labelled SH3 where the co-solute concentrations were incrementally increased up to 1 M (0, 0.05, 0.1, 0.25, 0.5, 1.0 M of IL/ salt concentration). Similarly, $^1H$-$^{15}N$ HSQC spectra were obtained for $Na_2SO_4$ and [Gdm]Cl salts at 0.4 M and 2.0 M concentration, respectively.

Combined $^1H$-$^{15}N$ chemical shift differences of the amide in 2D [$^1H$-$^{15}N$]-HSQC spectra, $\Delta\delta_{comb}$ or chemical shift perturbations (CSP), were calculated as

$$\Delta\delta_{\mathrm{comb}} = \sqrt{(\Delta\delta(\mathrm{H_N})^2 + (\alpha\Delta\delta\mathrm{N_H})^2} \qquad [S1]$$

where $\Delta\delta H_N$ and $\Delta\delta N_H$ are the $^1H$ and $^{15}N$ chemical shift differences of SH3 in the presence of added IL/ salt minus the same resonance in the absence of added species, normalized with the scaling factor α= 0.14 for most residues but α= 0.2 for glycine (10). To decide whether a given residue belongs to the class of interacting or non-interacting residues, we have calculated a cut-off value, based on the corrected standard deviation to zero, according to the method developed by (11). Since the total concentrations of ligand and protein are known during titration, $[L]_t$ and $[P]_t$, for residues above the cut-off, the $\Delta\delta_{comb}$ values were used to obtain the dissociation constant ($K_d$) from the titration experiments ([Ch][Glu] and [Bmim][dca]) by non-linear regression analysis according to ,

$$\Delta\delta_{\mathrm{comb}} = \Delta\delta_{\mathrm{max}} \frac{(\mathrm{K_d} + [L]_t + [P]_t) - \sqrt{(\mathrm{K_d} + [L]_t + [P]_t)^2 - (4[P]_t\,[L]_t)}}{2[P]_0} \qquad [S2]$$

where $\Delta\delta_{comb}$ is the combined chemical shift deviation from the free state defined by **Eq. S1** and $\Delta\delta_{max}$ is the maximum chemical shift change on saturation. Data are fitted to a single site binding model, using a least square fitting search of Microsoft Excel Solver to find the values of $K_d$ and the chemical shift of the fully saturated protein.

SH3 backbone assignments. Protein backbone resonance assignment experiments (Bruker standard pulse sequences) were recorded for 0.75 mM $^{15}N^{13}C$ drkN SH3 at 0, 0.35, 0.65, 1.0 M [Bmim][dca] and 1.0 M [Ch][Glu], comprising 2D $^{1}H$-$^{15}N$ HSQC and 3D triple resonance experiments HNCACB, HNcoCACB, HNCO, HNcaCO spectra. All spectra were collected using Echo/Antiecho-TPPI gradient selection, which was efficient at suppressing signals from ionic liquid. Acquisition parameters for 2D $^{1}H$-$^{15}N$ HSQC experiments were 2048 ($^{1}H$) and 512 ($^{15}N$) complex points for a SW of 12 ppm ($^{1}H$) and 32 ppm ($^{15}N$), with 16 scans. For 3D experiments, 2048 ($^{1}H$), 40 ($^{15}N$) and 256 ($^{13}C$) complex points for a SW of 12 ppm ($^{1}H$), 32 ppm ($^{15}N$) and 75 ppm for HNCAB/ HNcoCACB or 18 ppm for HNCO/ HNcaCO ($^{13}C$), were used with 16 scans except HNCACB with 32 scans. Non-uniform sampling (NUS) was used to optimize the resolution of the indirect dimensions in the available experiment time, where just 20-25% of the sparse data was recorded. The NUS acquired data were processed with SMILE algorithm within NMRPipe for spectra reconstruction (12). Sequential connectivities were performed with CcpNmr AnalysisAssign 3.0. (13) followed by manual verification. Backbone assignments for both folded and unfolded drkN SH3 in water were consistent with previous assignments under buffer conditions (8, 14), and tryptophan indole side chain resonances were assigned by analogy to previously published data (3). The backbone assignment for $U_{[Bmim][dca]}$ state led to an unambiguous assignment in the 2D $^{1}H$-$^{15}N$ HSQC even for very overlapped peaks.

The backbone $^{1}H$, $^{15}N$, $^{13}C\alpha$, $^{13}C\beta$, and $^{13}CO$ chemical shift assignments of drkN SH3 in water, 1.0 M [Ch][Glu] and 1.0 M [Bmim][dca] have been deposited in the BMRB database under the accession code 53285.

Secondary structure propensity (SSP). SSPs were calculated using $^{13}C_\alpha$ and $^{13}C_\beta$ chemical shifts of drkN SH3 as input according to the SSP protocol by (15). Positive SSP values ranging from 0 to 1 and negative values from 0 to -1 represent the propensities of α and β secondary structures, respectively. Reference values for random coil, secondary structure chemical shifts and standard deviations came from RefDB (16).

ZZex experiments. SH3 interconversion rates were determined by 2D longitudinal $^{15}N$ ZZex spectroscopy (17), using the pulse sequence provided by Bruker (hsqcetexf3gp). Sample conditions consisted of 1.1 mM protein in the absence or presence of 350 mM [ChGlu] or 150 mM [Bmim][dca]. A series of 13/14 2D exchange spectra were acquired at 293.2 K with variable mixing time ranging from 10 to 750 ms (10, 20, 40, 65, 100, 140, 180, 230, 320, 450, 550, 650, 750 ms of mixing time). Each 2D spectrum was recorded as a complex data matrix of 2048 ($^{1}H$) and 256 ($^{15}N$) points for a spectral width of 12 ppm ($^{1}H$) and 32 ppm ($^{15}N$), 32 scans per FID were obtained with a recycle delay of 1.2 s. These experiments were also employed to verify some of the assignments discussed above, and the ZZex data analysis is explained further below.

**NMR quantification of the F/U population.** The volume intensity for the two resonances from the same residue in each condition was extracted with CcpNmr 2.5. As the accuracy of the population determination is directly linked to the accuracy in peak volume determination, which, in turn, is dependent on the signal-to-noise ratio. We also calculated the volume intensity error for each peak volume using a Gaussian line shape fitting analysis by PINT (18, 19). The uncertainties in $\Delta G^{0'}{}_{u}$ values, as described in the main text, were propagated from peak volume error. The obtained $\Delta G^{0'}{}_{u}$ values were then fitted by a least squares analysis using Prism 8 (GraphPad Software). The *m*-value uncertainty is the standard error of the slope using a simple linear regression.

**NMR temperature scan analysis.** NMR temperature scan data were used to determine the population of folded ($p_f$) and unfolded ($p_u$) protein as a function of temperature, and from these populations, $\Delta G^{0'}{}_{u}$ was calculated as described in the main text. The data were fitted to the integrated Gibbs-Helmholtz19, assuming a constant heat capacity change upon unfolding ($\Delta C_p{}^{0}{}_{u}{}' = C_p{}^{0'}{}_{U} - C_p{}^{0'}{}_{F}$) over the temperature range studied (278 K to 313 K). Data were analyzed using Prism 8 (GraphPad Software) and the uncertainties are the standard deviation from the best-fit curves with nonlinear regression.

**Analysis of the ZZex spectroscopy data.** The time evolution of the resulting four peaks (auto-peaks (*ff* and *uu*) and exchange cross-peaks (*fu* and *uf*)) can be fit to the appropriate exchange model using the longitudinal Bloch-McConnell equations (21). Peak heights were used for *I(t)* values, where the full assignment was prepared with CcpNmr 2.5 and the integration was done with PINT (18, 19) using a Gaussian line shape fitting analysis, optimizing peak positions, line widths, and intensity. At *t*=0, the total intensity was set to the sum of the integrations of the *ff* and *uu* peaks and the cross-peak intensities were normalized to the nominal cross peak intensities. Non-linear least-squares and simultaneous fitting of $I_{ff}(t)$, $I_{uu}(t)$, $I_{fu}(t)$, $I_{uf}(t)$ curves for the measurement of chemical exchange ($k_{ex}$) and longitudinal $^{15}$N decay ($R_{1f}$ and $R_{1u}$) rates was conducted with a modified MATLAB script (22) in MATLAB R2017b (Mathworks) employing an exchange model for two-state interconversion as described by **Eqs. S3-6** that allow the extraction of $k_{fu}$ and $k_{uf}$ interconversion rates. The equations, for a two-state exchange process, describing the dependence of auto (*ff*, *uu*)- and cross (*fu, uf*) -peak heights on the variable mixing period, *t*, have been described before (17, 23, 24) and are given below:

$$I_{ff}(t) = \frac{1}{2}\left[\left(\frac{(1 - R^0_{1f} - R^0_{1u} + k_{ex}(p_u - p_f))}{(\lambda_+ - \lambda_-)}\right) e^{(-\lambda_- t)} + \left(\frac{(1 + R^0_{1f} - R^0_{1u} + k_{ex}(p_u - p_f))}{(\lambda_+ - \lambda_-)}\right) e^{(-\lambda_+ t)}\right] \quad [S3]$$

$$I_{uu}(t) = \frac{1}{2}\left[\left(\frac{(1 + R^0_{1f} - R^0_{1u} + k_{ex}(p_u - p_f))}{(\lambda_+ - \lambda_-)}\right) e^{(-\lambda_- t)} + \left(\frac{(1 - R^0_{1f} - R^0_{1u} + k_{ex}(p_u - p_f))}{(\lambda_+ - \lambda_-)}\right) e^{(-\lambda_+ t)}\right] \quad [S4]$$

$$I_{fu}(t) = \frac{k_{ex}p_f}{(\lambda_+ - \lambda_-)}\left(e^{(-\lambda_- t)} - e^{(-\lambda_+ t)}\right) \quad [S5]$$

$$I_{uf}(t) = \frac{k_{ex}p_u}{(\lambda_+ - \lambda_-)}\left(e^{(-\lambda_- t)} - e^{(-\lambda_+ t)}\right) \quad [S6]$$

*where I* refers to the time dependence of the transfer amplitudes (build-up curves) for the *ff*, *uu, f* to *u (fu)* and *u* to *f (uf)* interconversions. *p* refers to the fractional population of the indicated state, $k_{ex}$ is the stochastic exchange of molecules between the two states per second (s), *t* is time in s. $R_{1f}$ and $R_{1u}$ are the longitudinal relaxation rate constants of magnetization in sites *f* and *u*, respectively. $I_{ff}(0)$ and $I_{uu}(0)$ denote the amount of longitudinal nitrogen magnetization associated with the folded and unfolded states at the start of the mixing period t. In the limit, $R_{1f} = R_{1u}$ and $k_{fu} = k_{uf}$. $\lambda_\pm$ denotes the eigenvalues of the 2x2 dynamics matrix ($a_{1,2}$) describing the loss of magnetization in the folded and unfolded states due to longitudinal relaxation and chemical exchange, given by,

$$\lambda_\pm = \frac{1}{2}\left\{R^0_{1f} + R^0_{1u} + k_{ex} \pm \left[\left(R^0_{1f} + R^0_{1u} + k_{ex}\,(p_u - p_f)\right)^2 + 4p_f p_u {k_{ex}}^2\right]\right\} \quad [S7]$$

Where the intensities of the auto-peaks decrease with increasing *t*, while the exchange peaks increase to a maximum in intensity (at $t = \frac{ln\left(\frac{\lambda_-}{\lambda_+}\right)}{(\lambda_- - \lambda_+)}$) and subsequently decrease in intensity.

<u>Analysis of the folding thermodynamics and kinetics.</u> A two-site exchange process is considered according to:

$$F \underset{k_f}{\overset{k_u}{\rightleftarrows}} U \quad [S8]$$

where the chemical shift difference between sites is δ*ω*, the equilibrium populations of states *F* and *U* are $p_f$ and $p_u$ with $p_f + p_u = 1$. The exchange first order rates for the magnetization converting from site f to *u* ($k_u$) – unfolding rate constant, and *u* to *f* ($k_f$) – folding rate constant*,* were calculated using the relative populations of the two states taken from integrations:

$$k_{ex} = k_u + k_f = \frac{k_u}{p_u} = \frac{k_f}{p_f} \quad [S9]$$

$$k_u = k_{ex}p_u \quad [S10]$$

$$k_f = k_{ex}p_f \quad [S11]$$

The unfolding free energy (F→U, $\Delta G^{0'}{}_{u}$) per residue can be calculated using the following relation:

$$\Delta G_u^{0\prime} = -RTln\left(\frac{k_u}{k_f}\right) \qquad [S12]$$

The (un)folding rates extracted from ZZex data allow the characterization of how the barriers that define SH3 folding (U→TS‡) and unfolding (F→TS‡) are modified with ILs. Based on the transition-state theory (25), the free energies required to reach the transition state (TS‡) from the unfolded ensemble, $\Delta G_{U\rightarrow TS\ddagger}{}^{0'\ddagger}$, and the folded state, $\Delta G_{F\rightarrow TS\ddagger}{}^{0'\ddagger}$, are determined by the Eyring-Polanyi equation (26, 27),

$$\Delta G^{0'\ddagger}_{F,U\rightarrow TS\ddagger} = -RTln\left(\frac{k_{F,U\rightarrow TS\ddagger}h}{k_B T}\right) \qquad [S13]$$

where $\Delta G^{0'\ddagger}{}_{F,U\rightarrow TS\ddagger}$ is the modified standard-state activation free energy for folding or unfolding at absolute temperature $T$, $R$ is the gas constant, $k$ is the folding or unfolding rate at T, $h$ is Planck's constant, and $k_B$ is the Boltzmann constant. For clarity, $k_u = k_{F\rightarrow TS\ddagger}$, $k_f = k_{U\rightarrow TS\ddagger}$, and $\Delta G^{0'}{}_{F\rightarrow U} = \Delta G^{0'}{}_{u}$.

The viscosity effects were also considered, since (28) shows that drkN SH3 folding is linearly dependent on solvent viscosity. In this context, Kramers' model for diffusive barrier crossing (29) appears to be more appropriate than the original transition state formalism because protein folding necessarily involves diffusional events, as the expanded and hydrated unfolded state polypeptide chain collapses toward a more compact and conformationally restricted folded protein (water is expelled from the interior of the protein). In cases where the rate-limiting step involves such diffusive processes and based on the Stokes law, a 1/η dependence of $k_{fu}$ and $k_{uf}$ on solvent viscosity is expected, whereas rate constants will be independent of solvent viscosity if the rate-limiting step involves only rearrangements that are limited by the internal friction of the protein:

$$k_{F,U\rightarrow TS\ddagger}(T,\eta) = \frac{A}{\eta} exp\left(\frac{-\Delta G^{0'\ddagger}_{F,U\rightarrow TS\ddagger}}{k_B T}\right) \qquad [S14]$$

where A is a temperature and viscosity-independent constant. Thus, to separate the viscosity effects from that result from increased stability (i.e., viscosity does not change the equilibrium of the folding reaction but affects both folding and unfolding to the same extent), we adjusted the folding/unfolding rates accordingly (30, 31):

$$k_{0\,(F,U\rightarrow TS\ddagger)} = k_{F,U\rightarrow TS\ddagger}\left(\frac{\eta_c}{\eta_{water}}\right) = k_{F,U\rightarrow TS\ddagger}\eta_{rel} \qquad [S15]$$

where $k_0$ is the viscosity-corrected rate, $k$ is the folding/unfolding rate before viscosity correction, $\eta_c$ is the viscosity of the solution, $\eta_{water}$ is the viscosity of pure water at 293.2 K (1.002 cP) (32), and $\eta_{rel}$ is the viscosity corrected to water at 293.2 K. Viscosity measurements and extrapolations are described below. Non- and viscosity-corrected rates were listed for the different conditions (water, 0.35 M [Ch][Glu] and 0.15 M [Bmim][dca] at 293.2 K) in **Table S5**.

Rearranging **equation S15**, the free energies per residue required to reach the transition state (TS‡) are determined by:

$$\left(\Delta G^{0'\ddagger}_{F,U\rightarrow TS\ddagger}\right)_{corr} = -RTln\left(\frac{k_{0\,(F,U\rightarrow TS\ddagger)}h}{k_B T}\right) \qquad [S16]$$

Together, $(\Delta G_{F\rightarrow TS\ddagger}{}^{0'\ddagger})_{corr}$ and $(\Delta G_{U\rightarrow TS\ddagger}{}^{0'\ddagger})_{corr}$ describe the correct equilibrium between the thermodynamic states and the transition state:

$$\Delta G_u^{0\prime} = \left(\Delta G^{0'\ddagger}_{F\rightarrow TS\ddagger}\right)_{corr} - \left(\Delta G^{0'\ddagger}_{U\rightarrow TS\ddagger}\right)_{corr} \qquad [S17]$$

The average of residue-specific values of $k_u$, $k_f$, and other extracted parameters was employed in all analyses, and errors were estimated from the standard deviation of the values over the residues that were included in the average.

**Viscosity measurements and extrapolations.** Viscosity of pure water at 293.2 K is 1.002 cP(32). The relative viscosity ($\eta_{rel}$) to water at 293.2 K determined for 0.35 M [ChGlu] and 0.15 M [Bmim][dca] are 1.39 and 1.02, respectively. The details of the measurements and extrapolations are as follows.

0.35 M [Ch][Glu] viscosity solution. Viscosities of 0.1 M, 0.5 M, 1.0, and 1.5 M aqueous-[Ch][Glu] were measured from 294.2 to 312.2 K in 2 K increments at atmospheric pressure using an automated SVM 3000 rotational Stabinger viscometer–densimeter (Anton Paar). The SVM 3000 uses Peltier elements for fast and efficient thermal stability. At each concentration, to estimate an uncertainty from standard deviation, a minimum of three viscosity values were measured for three temperatures (298.2, 304.2, and 310.2 K). **Table S8** shows the measured viscosities in triplicate for each [Ch][Glu] concentration for three temperatures. According to these measurements, the viscosity as a function of $T$ was fit to an exponential function, which was used to extrapolate the $\eta$ at 293.2 K. Likewise, the dependency of the solvent viscosity as a function of [Ch][Glu] concentration ([Ch][Glu]) at 293.2 K can be described by an empirical exponential equation:

$$\eta\ (293.2\ K, [Ch][Glu]) = 1.046677 \exp(0.809516\ [Ch][Glu]) \qquad [S18]$$

This equation has no physical meaning but describes the dependence of viscosity on [Ch][Glu] concentration very well at 293.2 K. Viscosity of [Ch][Glu] IL 0.35 M at 293.2 K was calculated to 1.390 cP.

0.15 M [Bmim][dca] viscosity solution. Viscosity of aqueous-[Bmim][dca] IL 0.15 M at 293.2 K was determined using **Eq. S19** based upon Eyring's theory and a modified two-suffix-margules excess Gibbs energy model (Eyring-MTSM) to correlate the dynamic viscosities of binary mixtures of ILs with water (33):

$$\ln(\eta_{mix}) = x_1 ln(\eta_1) + x_2 ln(\eta_2) + \alpha_{12}\frac{x_1 x_2 G_{12} G_{21}}{(x_1 G_{12} + x_1)(x_1 G_{12} + x_1)} \qquad [S19]$$

with

$$G_{12} = \exp\left(\frac{\tau_{12}}{RT}\right), \qquad \tau_{12} = g_{12} - g_{22} \qquad [S20]$$

$$G_{21} = \exp\left(\frac{\tau_{21}}{RT}\right), \qquad \tau_{21} = g_{21} - g_{11} \qquad [S21]$$

and

$$\alpha_{12} = \alpha_{12}^{(0)} + \frac{\alpha_{12}^{(1)}}{T} \qquad [S22]$$

where $x_1$, $x_2$, $\eta_1$, $\eta_2$ are the mole fraction and viscosities of component 1 and 2, respectively, $g_{ij}$ is the potential energy between components i and j, R is the gas constant, α12, τ12 and τ21 are adjustable parameters, and $\alpha_{12}(0)$ and $\alpha_{12}(1)$ are adjustable temperature dependent considerations. The interaction parameters of the model for [Bmim][dca] (component 1) and water (component 2) were taken from (34), $\alpha_{12}(0)$, $\alpha_{12}(1)$, τ12 and $\tau_{21}$ are 0.9838, 428.2, 407.3, -467.9, respectively. [Bmim][dca] pure viscosity ($\eta_1$) is 39.14 cP at 293.2 K (35). The viscosity of aqueous-[Bmim][dca] IL 0.15 M at 293.2 K was calculated to 1.018 cP.

#### SI Figures

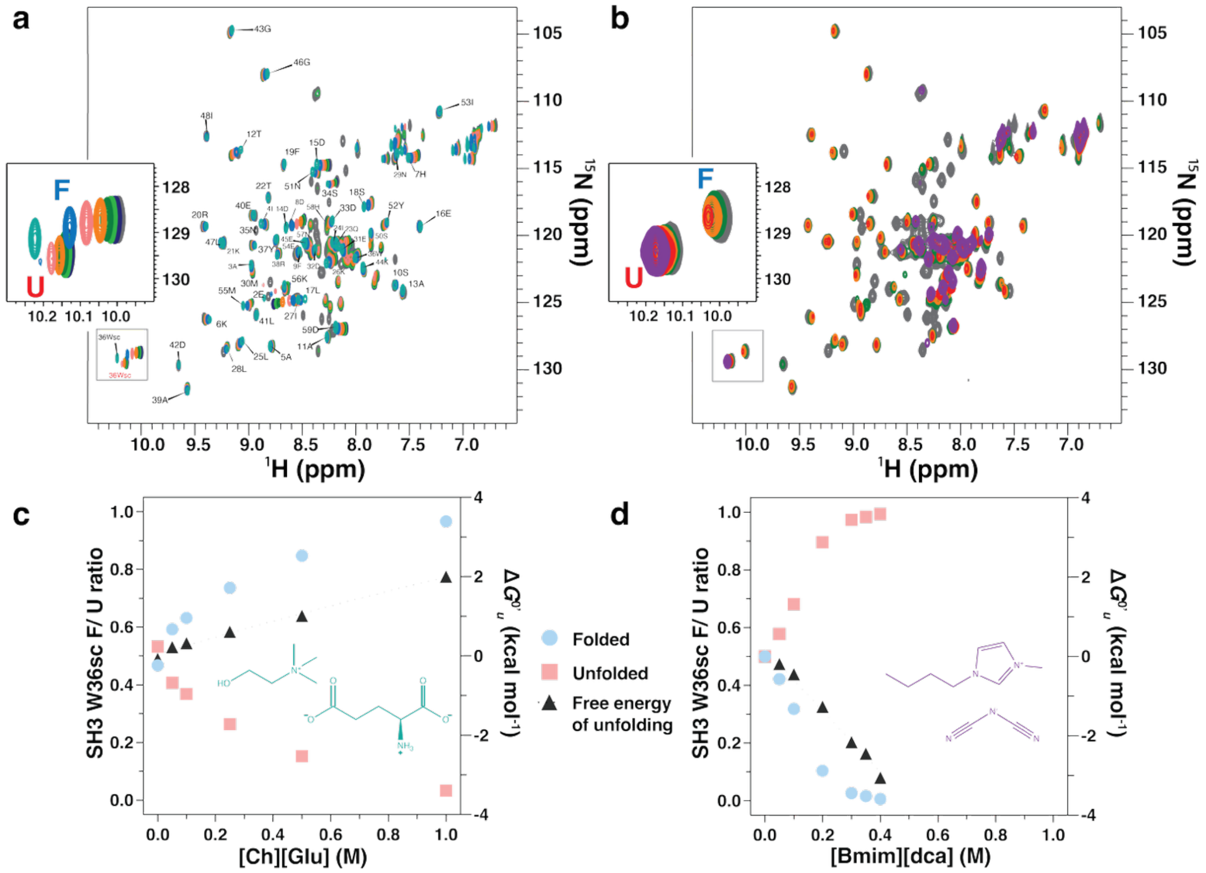

**Fig. S1: [$^{1}$H,$^{15}$N]-HSQC titration with [Ch][Glu] and [Bmim][dca] and their corresponding folded/ unfolded populations.** Overlay of 2D $^{1}$H-$^{15}$N HSQC spectra of drkN SH3 acquired in (**a**) [Ch][Glu] titration (0, 0.01, 0.025, 0.05, 0.1, 0.25, 0.5 and 1.0 M) or in (**b**) [Bmim][dca] titration (0, 0.05, 0.1, 0.2 and 0.3 M). The label shows the assignment for the folded state, and the inlay shows the indole W36 side chain peaks of the folded (F) and unfolded (U) states. All the spectra were acquired in water, to avoid interference from the buffer, solution pH ≈ 7.1, at 298.2 K and 600.13 MHz. (**c**, **d**) Populations and Gibbs free energy of unfolding ($\Delta G^{0'}{}_{u}$) were calculated as a function of (c) [Ch][Glu] and (d) [Bmim][dca] concentration. Based on each $^{1}$H-$^{15}$N HSQC spectrum, peak volume intensities of the exposed indole side chain of W36 were used to calculate the populations of the folded state ($p_f$, blue) and the unfolded ensemble ($p_u$, red). For each titration point, the Gibbs free energy of unfolding ($\Delta G^{0'}{}_{u}$, black) was calculated assuming a two-state unfolding model.

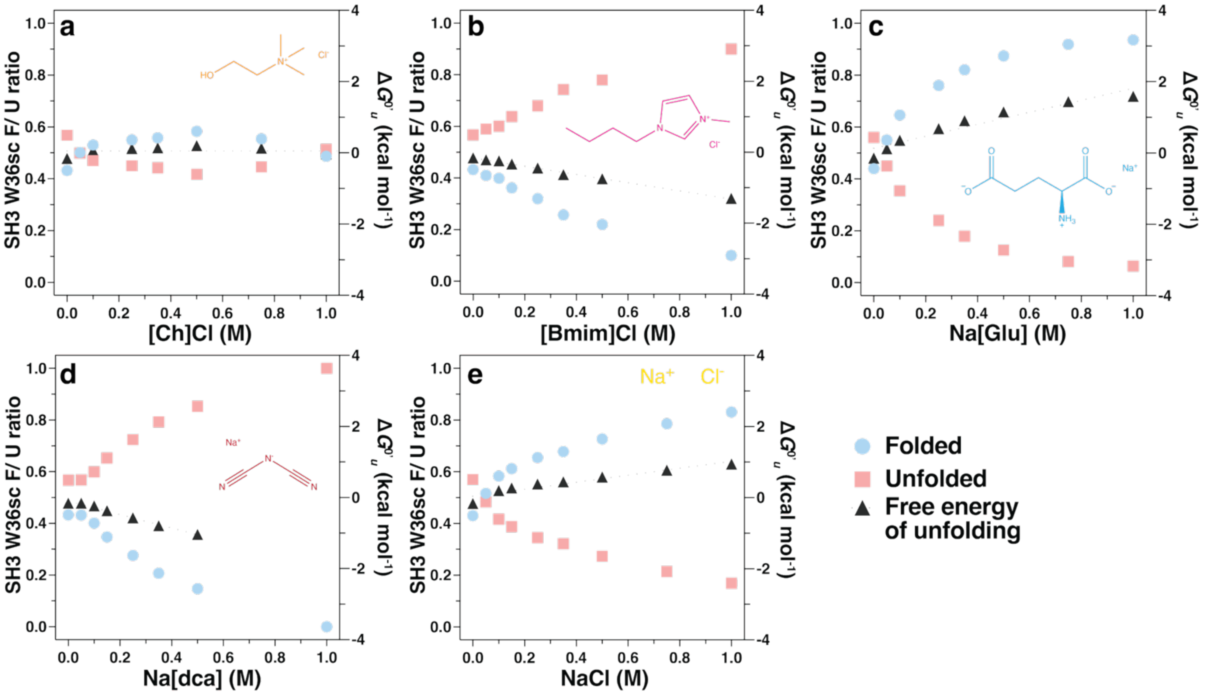

**Fig. S2: Populations of folded and unfolded states along salt titrations.** Populations and Gibbs free energy of unfolding ($\Delta G^{0'}{}_{u}$) were calculated as function of salts concentration related with [Ch][Glu] and [Bmim][dca: (**a**) [Ch]Cl, (b) [Bmim]Cl, (**c**) Na[Glu], (**d**) Na[dca]; and (**e**) NaCl as a reference. Based on each $^{1}$H-$^{15}$N HSQC spectrum, peak volume intensities of the exposed indole side chain of W36 were used to calculate the populations of the folded state ($p_f$, blue) and the unfolded ensemble ($p_u$, red). For each titration point, the Gibbs free energy of unfolding ($\Delta G^{0'}{}_{u}$, black) was calculated assuming a two-state unfolding model.

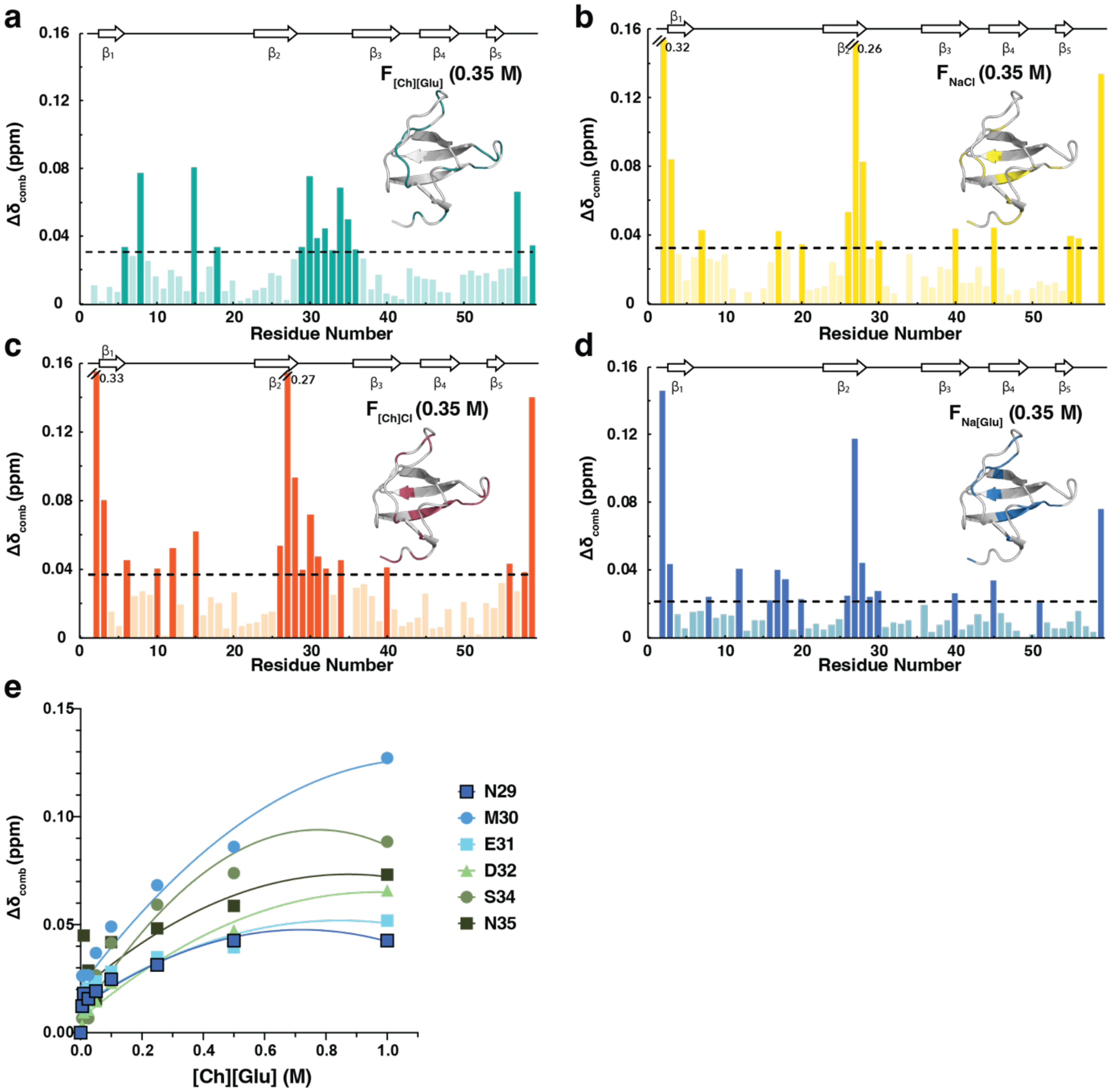

**Fig. S3: [Ch][Glu] IL and salts–protein interactions in the folded state.** (**a**) [Ch][Glu], (**b**) NaCl, (**c**) [Ch]Cl, (**d**) Na[Glu]. In (b) and (c), the magnitude of the CSPs of residues Q2 and L28 is higher than 0.16; therefore, to maintain the same plot scale (for easier comparison), the bars corresponding to these residues were cut, and their magnitude value is written at their side. The residues that show a combined chemical shift above the threshold (dashed line) are colored in a darker tone. Above each plot, the secondary structure of the protein is depicted. The combined chemical shifts were calculated in the presence of [Ch][Glu] or respective salts at a concentration of 0.35 M and against the folded chemical shifts in water ($F_{water}$). (e) $\Delta\delta_{comb}$ versus [Ch][Glu] for residues N29–N35.

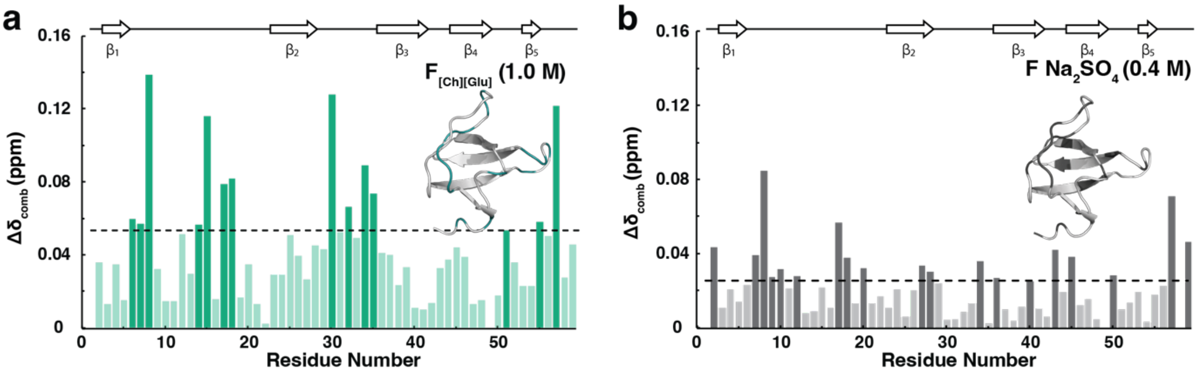

**Fig. S4: Combined chemical shift of the folded drkN SH3 in the presence of stabilizer IL or salt.** Combined chemical shift of the folded drkN SH3 in the presence of (**a**) 1.0 M of [Ch][Glu] (purple) and (**b**) 0.40 M of $Na_2SO_4$ (grey). The dashed lines correspond to the cut-off value determined for both cosolutes and are colored accordingly. Above the plot, the secondary structure of the protein is depicted. The combined chemical shifts were calculated against the folded chemical shifts in water ($F_{water}$).

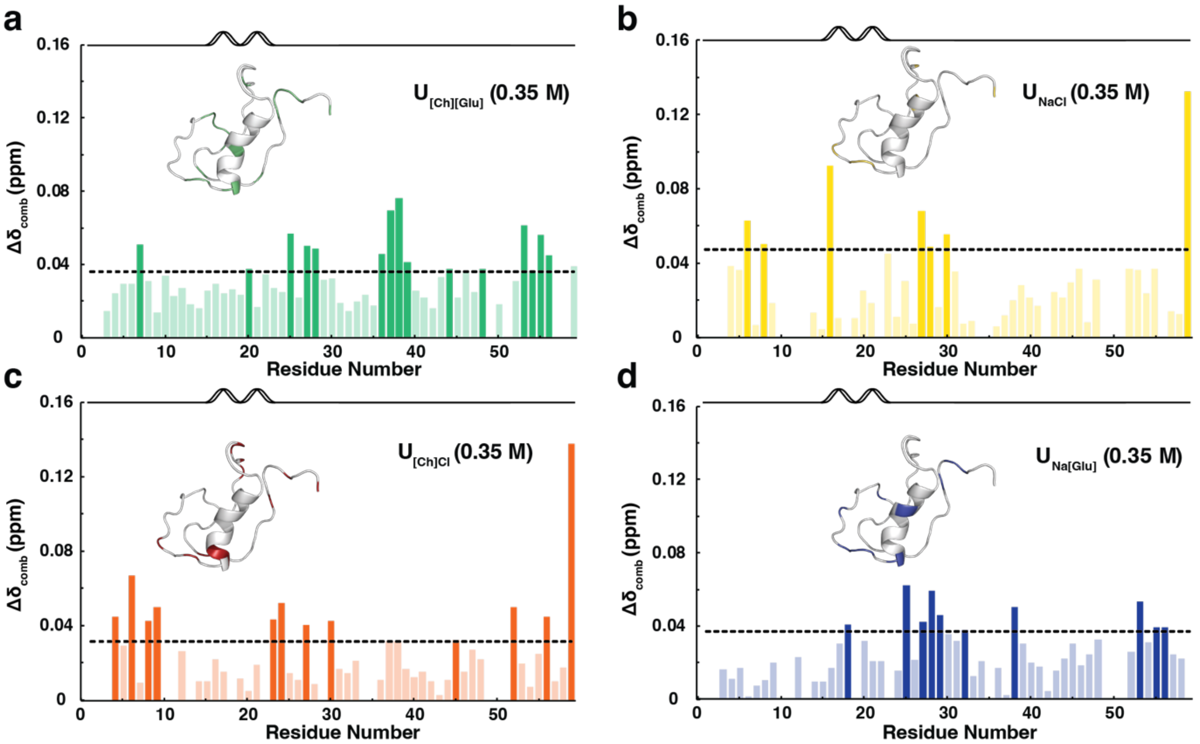

**Fig. S5: [Ch][Glu]/ionic salts–protein interactions in the unfolded state.** (**a**) [Ch][Glu], (**b**) NaCl, (**c**) [Ch]Cl, (**d**) Na[Glu]. The residues that show a combined chemical shift above the threshold (dashed line) are colored in a darker tone. Above the plot, the predicted secondary structure of the protein is depicted. The combined chemical shifts were calculated in the presence of 0.35 M of [Ch][Glu] and respective salts and against the unfolded chemical shifts in water ($U_{water}$).

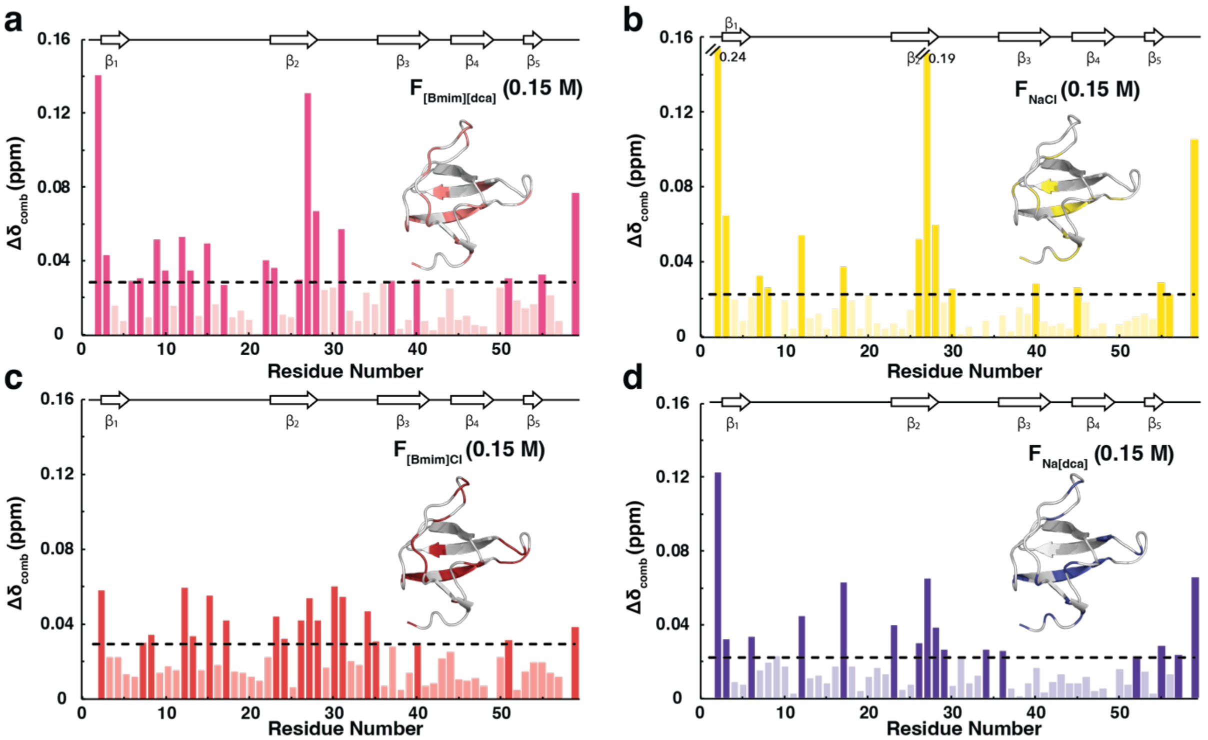

**Fig. S6: [Bmim][dca]/ionic salts–protein interactions in the folded state.** (**a**) [Bmim][dca], (**b**) NaCl, (**c**) [Bmim]Cl, (**d**) Na[dca]. In (b), the magnitude of the CSPs of residues Gln2 and Leu28 is higher than 0.16; therefore, to maintain the same plot scale (for easier comparison), the bars corresponding to these residues were cut, and their magnitude value is written at their side. The residues that show a combined chemical shift above the threshold (dashed line) are colored in a darker tone. Above each plot, the secondary structure of the protein is depicted. The combined chemical shifts were calculated in the presence of 0.15 M of [Bmim][dca] and respective salts and against the folded chemical shifts in water ($F_{water}$).

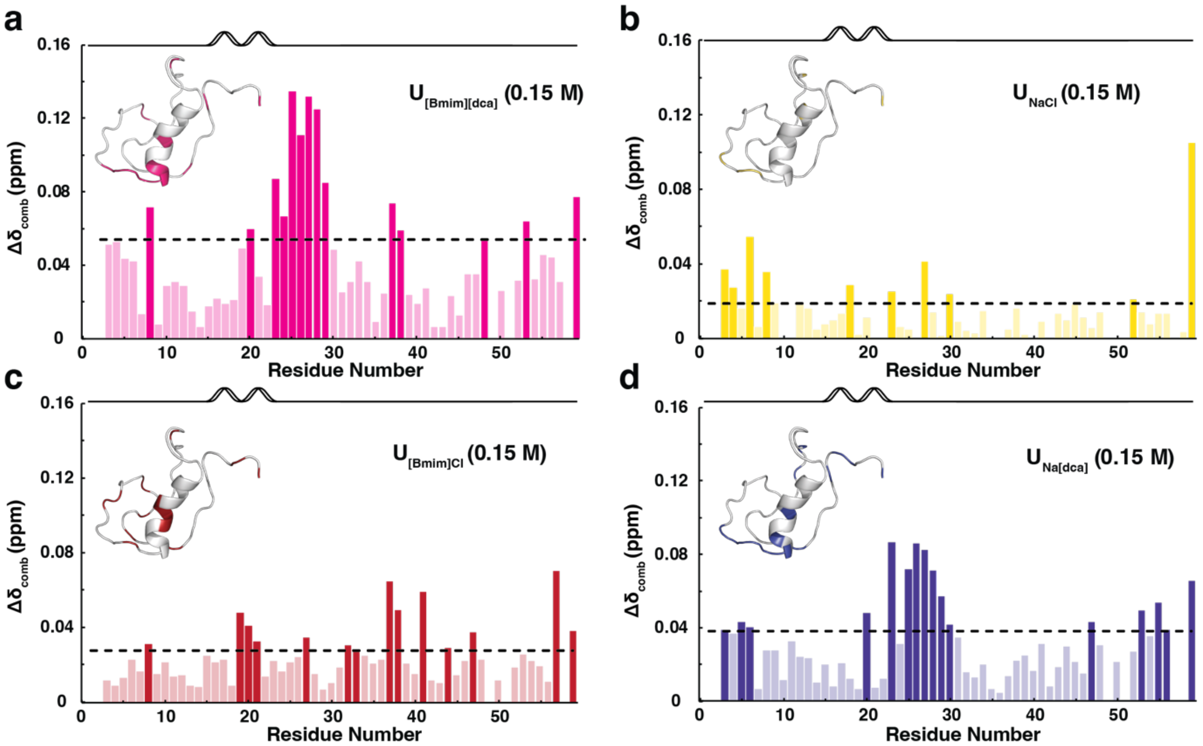

**Fig. S7: [Bmim][dca]/ionic salts–protein interactions in the unfolded state.** (**a**) [Bmim][dca], (**b**) NaCl, (**c**) [Bmim]Cl, (**d**) Na[dca]. The residues that show a combined chemical shift above the threshold (dashed line) are colored in a darker tone. Above the plot, the predicted secondary structure of the protein is depicted. The combined chemical shifts were calculated in the presence of 0.15 M of [Bmim][dca] and respective salts and against the unfolded chemical shifts in water ($U_{water}$).

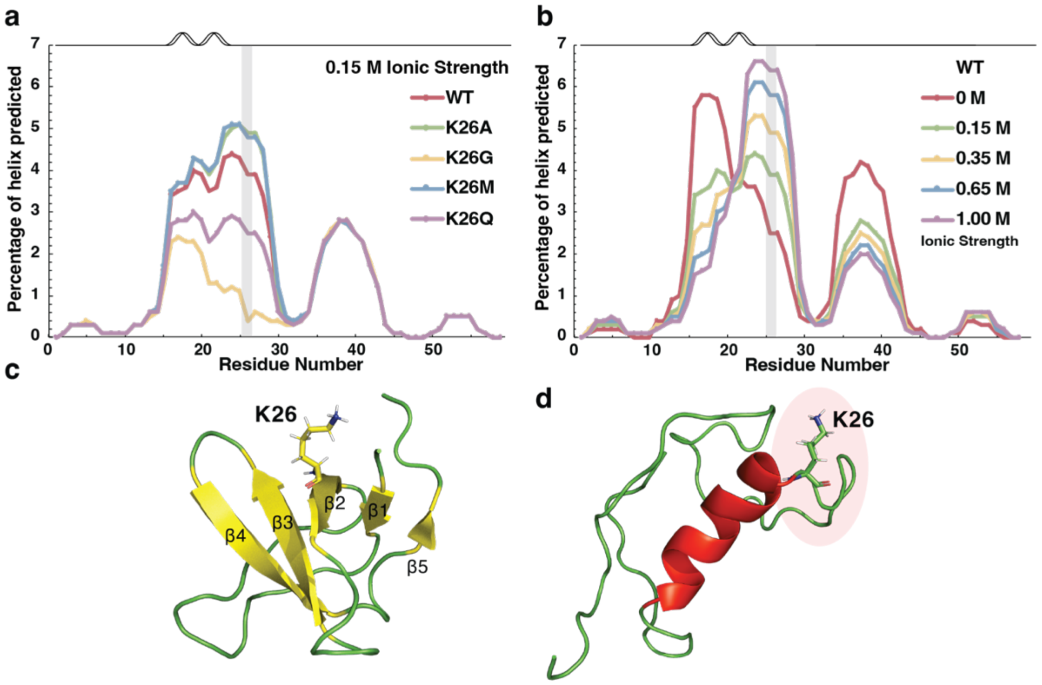

**Fig. S8: AGADIR prediction of the fractional α-helical population of drkN SH3 domain as a function of residue.** (**a**) Wild-type (WT - red) versus mutants (K26A – green; K26G – yellow; K26M – blue and K26Q – purple); ionic strength 0.15 M. (**b**) Wild-type at different ionic strengths (0 M – red; 0.15 M– green; 0.35 M – yellow; 0.65 M – blue and 1.0 M – purple). The predictions were performed at 298 K, pH = 7. Above the plot, the predicted secondary structure of the protein is depicted. Residue 26 is highlighted by a grey box. 3D structure for the (c) folded state (PDB: 2A36) and (d) representative of the unfolded ensemble in water (PED00022, #576), the K26 is highlighted by sticks.

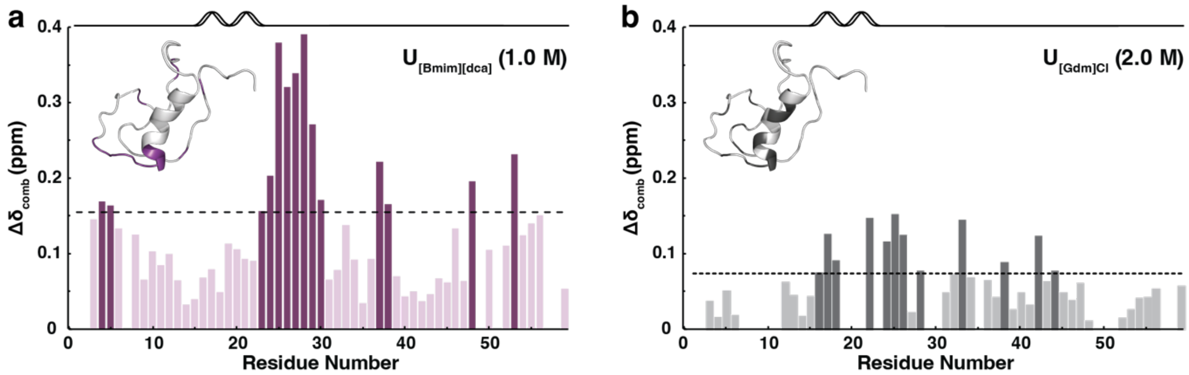

**Fig. S9: CSP of the unfolded drkN SH3 in the presence of [Bmim][dca] and [Gdm]Cl.** Combined chemical shift of the unfolded drkN SH3 in the presence of (**a**) 1.0 M [Bmim][dca] (purple) and (**b**) 2.0 M [Gdm]Cl (dark purple). The dashed lines correspond to the cut-off value determined for both cosolutes and are colored accordingly. Above the plot, the predicted secondary structure of the protein is depicted. The combined chemical shifts were calculated against the unfolded chemical shifts in water ($U_{water}$).

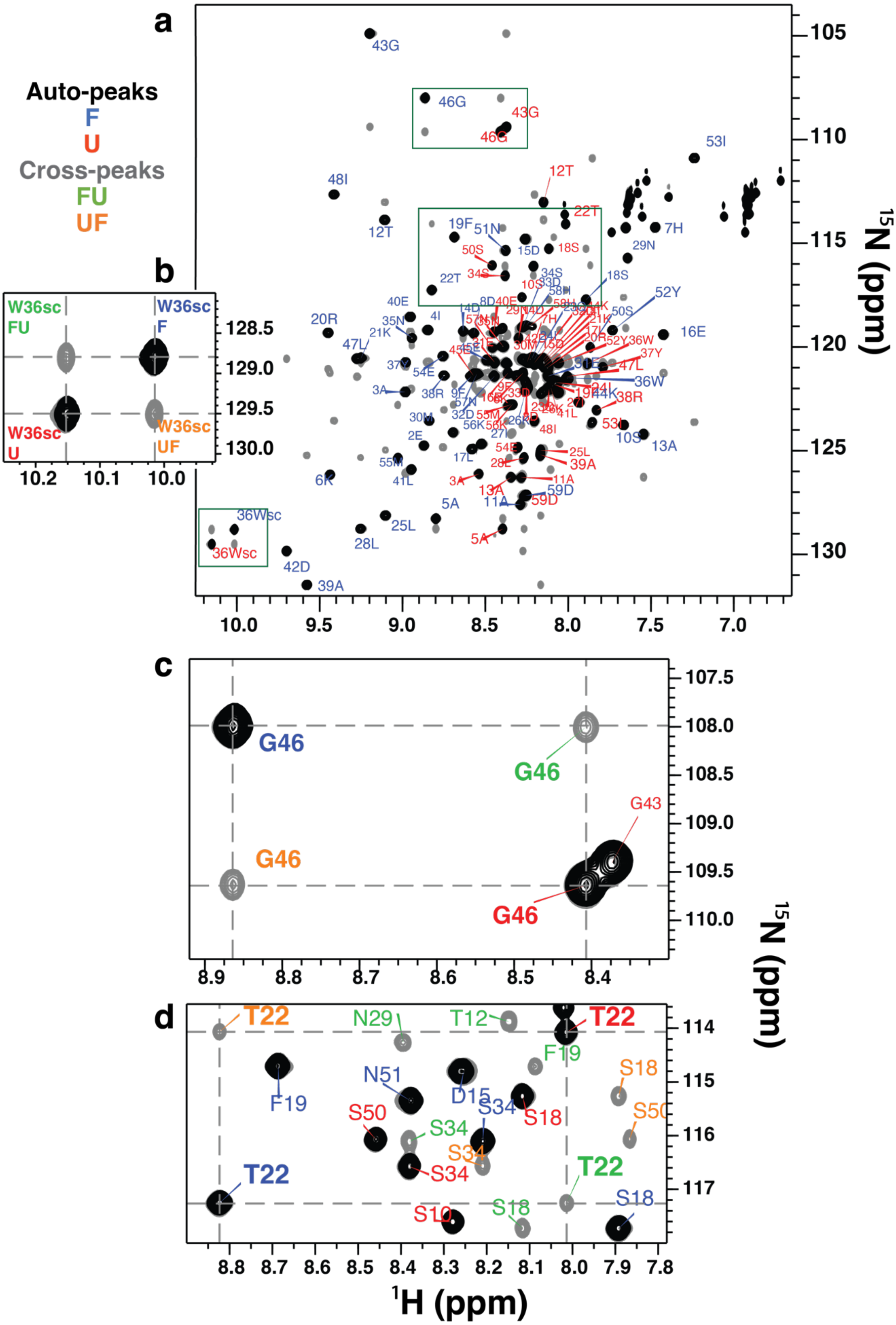

**Fig. S10: $^{1}H$-$^{15}N$ ZZ-exchange spectrum of drkN SH3 in water.** The spectrum was acquired with 320 ms of mixing time at 600.13 MHz, 293.2 K for 1.1 mM of $^{15}N$-labelled drkN SH3 in water. (**a**) full spectrum. The different inlays show (**b**) side chain of W36, (**c**) G46, and (**d**) T22 NH auto- and cross-peaks. Peaks for the folded and unfolded forms of the residue (black) are labelled with F (blue) and U (red), respectively, and dotted lines indicate the exchange cross-peaks (gray), FU (green), and UF (orange).

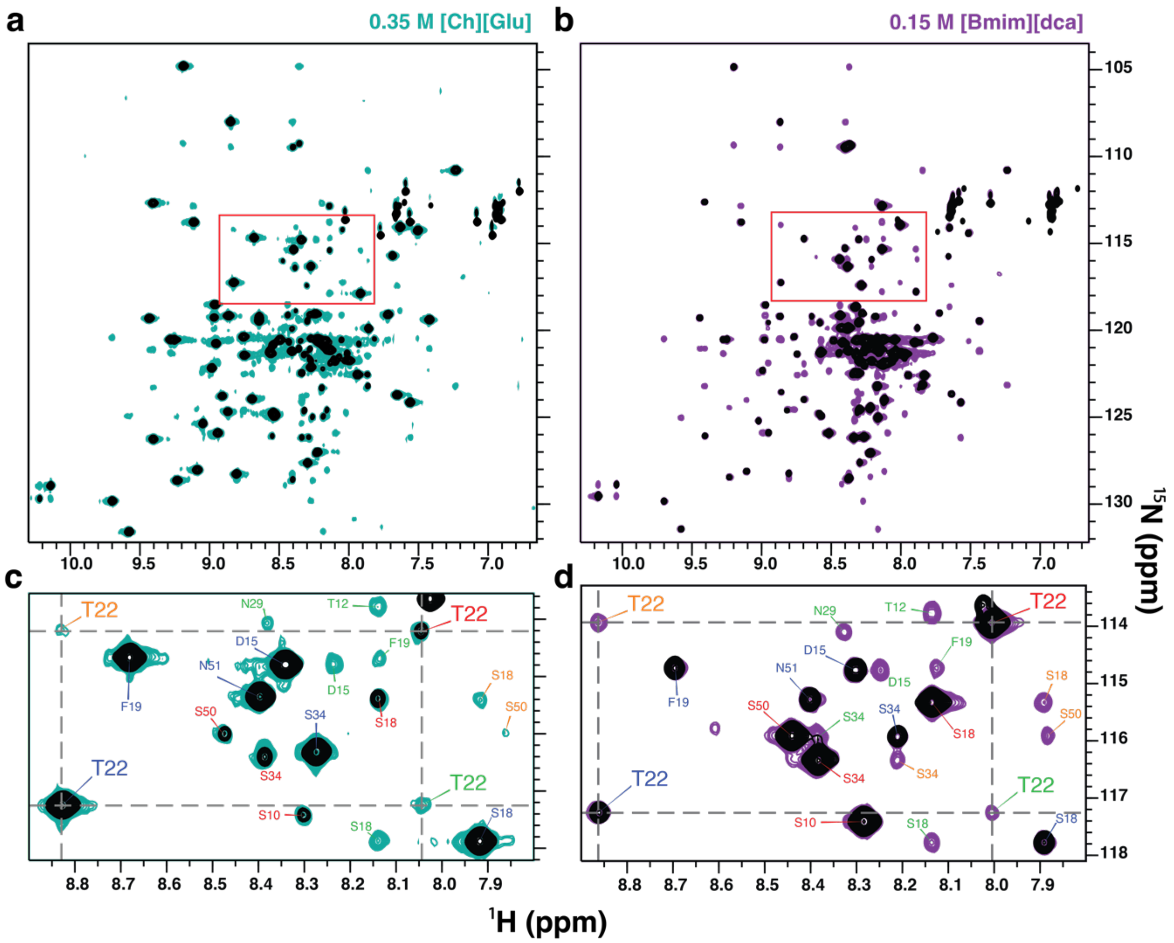

**Fig. S11: $^{1}$H-$^{15}$N ZZ-exchange spectra of drkN SH3 in 0.35 M [Ch][Glu] and 0.15 M [Bmim][dca].** The spectra were acquired with 320 ms of mixing time at 600.13 MHz, 293.2 K for 1.1 mM of $^{15}$N labelled drkN SH3 in (**a**) 0.35 M [Ch][Glu] and (**b**) 0.15 M [Bmim][dca]. The inlay (**c,d**) shows T22 NH auto- and cross-peaks. Peaks for the folded and unfolded forms of the residue (black) are labelled with F (blue) and U (red), respectively, and dotted lines indicate the exchange cross-peaks (gray), FU (green), and UF (orange).

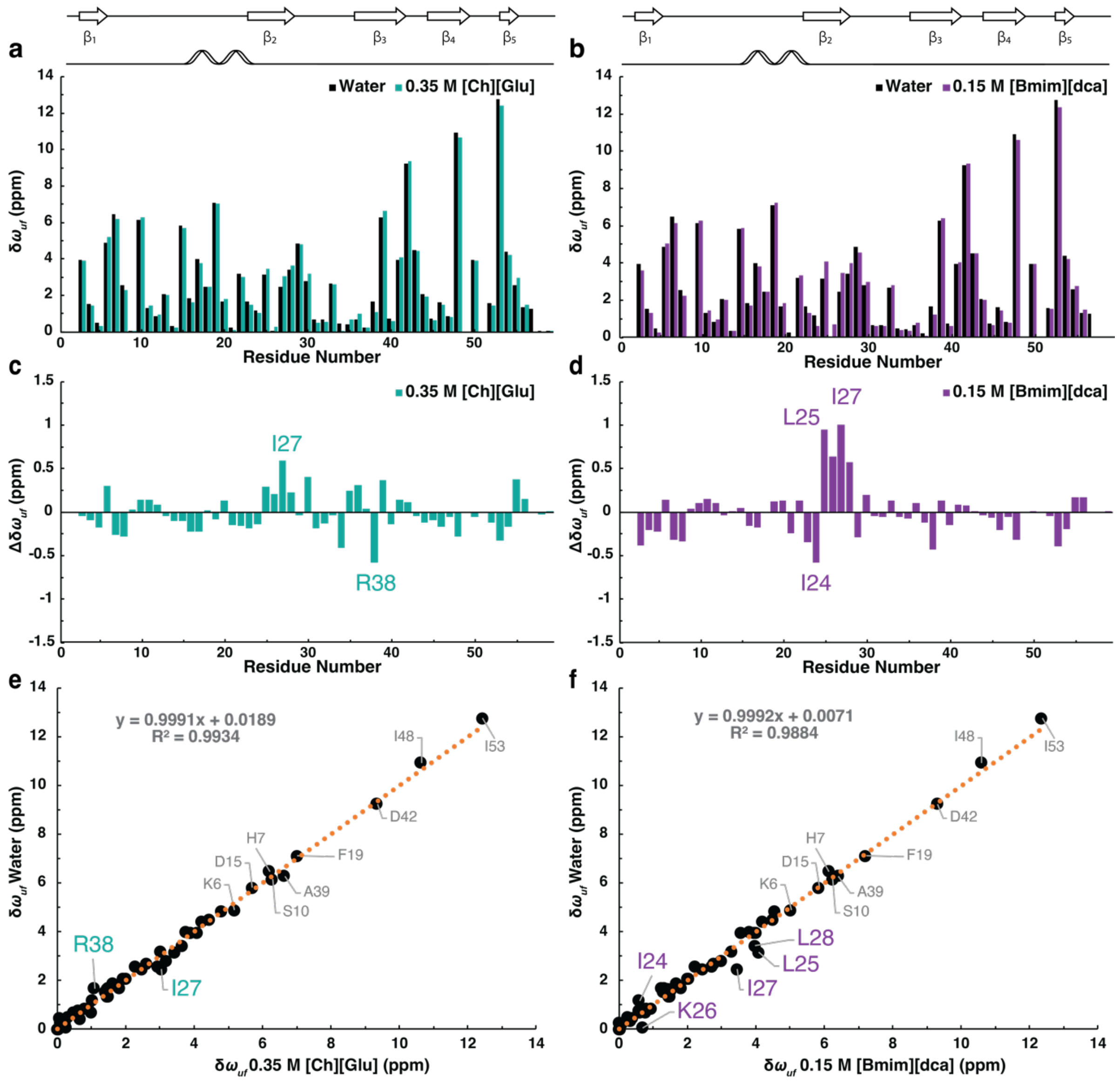

**Fig. S12: Correlation of $\delta\omega_{UF}$ for water, [Ch][Glu] and [Bmim][dca] ILs.** $^{15}N$ chemical shift difference between $F_{exch}$ and $U_{exch}$, $\delta\omega_{UF}$, extracted directly from ZZex-spectra for water (gray) and (**a**) 0.35 M [Ch][Glu] or (**b**) [Bmim][dca] (black). The difference, $\Delta\delta\omega_{UF} = \delta\omega_{UF,\ cosolute} - \delta\omega_{UF,\ water}$, as calculated for (**c**) [Ch][Glu] and (**d**) [Bmim][dca]. Correlation between $^{15}N$ chemical shift difference between $F_{exch}$ and $U_{exch}$, $\delta\omega_{UF}$, obtained directly from ZZex-spectra for water and (**e**) 0.35 M [Ch][Glu] or (f) [Bmim][dca]. The gray label indicates $\delta\omega_{UF}$ residues > 5 ppm, and the purple/orange label indicates the deviations ($|\Delta\delta\omega_{UF} = \delta\omega_{UF}(IL) - \delta\omega_{UF}\ (water)| > 0.5$ ppm).

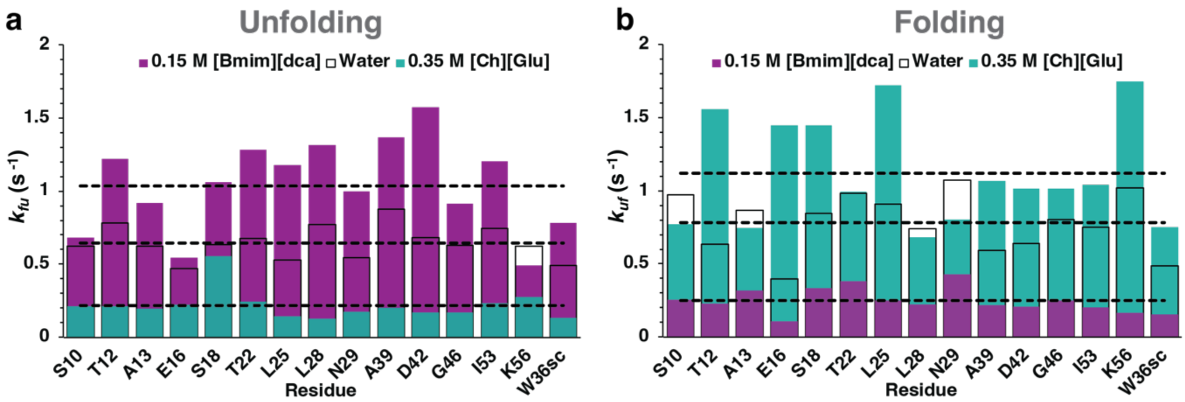

**Fig. S13: Viscosity-corrected rates for (un)folding interconversion of SH3 in water and ILs.** Interconversion rates for (**a**) unfolding and (**b**) folding ($k_{fu}$ and $k_{uf}$, respectively) as determined from the fitting of experimental ZZex data in water, 0.35 M [Ch][Glu], and 0.15 M [Bmim][dca]. These rates were corrected accordingly with their relative viscosities ($\eta_{rel}$ used in 0.35 M [ChGlu], and 0.15 M [Bmim][dca] are 1.39, and 1.02, respectively.) Black, turquoise, and purple bars correspond to the viscosity-corrected rate in water, [Ch][Glu], and [Bmim][dca], respectively. The horizontal dashed lines correspond to their 15-residue average.

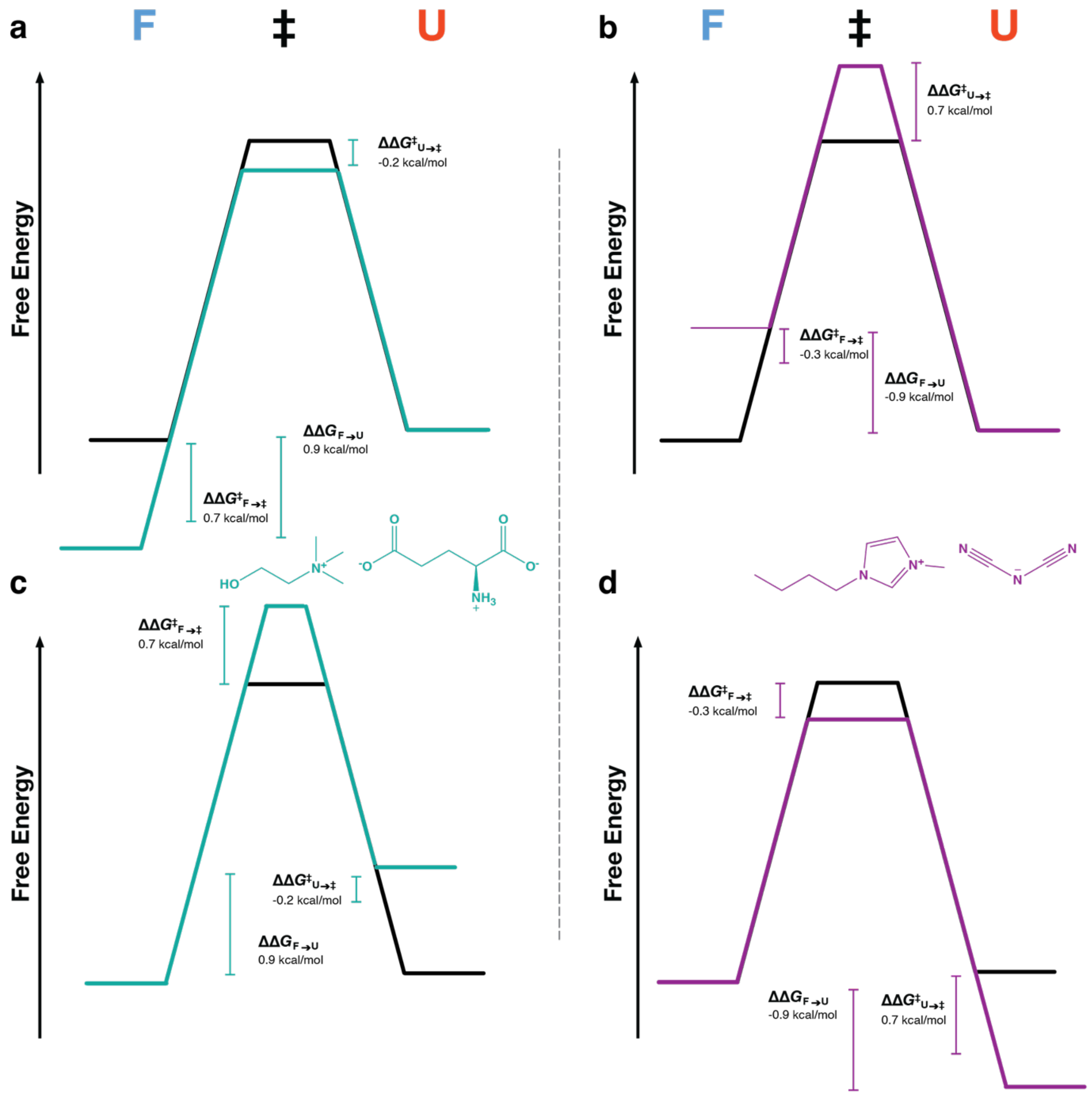

**Fig. S14: ILs dependence of the drkN SH3 free energy landscape with changes in the transition state.** (**a,c**) free energy landscape model for the stabilization of SH3 in [Ch][Glu]. (**b,d**) free energy landscape model for the destabilization of SH3 in [Bmim][dca]. Model (a,b) assumes that the unfolded state energy is unaffected by ionic liquid and that the thermodynamic and kinetic behavior can be explained by changes in the energies of the folding transition and folded states. Model (c,d) assumes that the folded state energy is unaffected by ionic liquid and that the thermodynamic and kinetic behavior can be explained by changes in the energies of the folding transition and unfolded states. $\Delta\Delta G^{0'}{}_{u}$, $\Delta\Delta G^{0'\ddagger}{}_{F,U\rightarrow TS\ddagger}$ average values and uncertainties were calculated from the equations described in the main text.

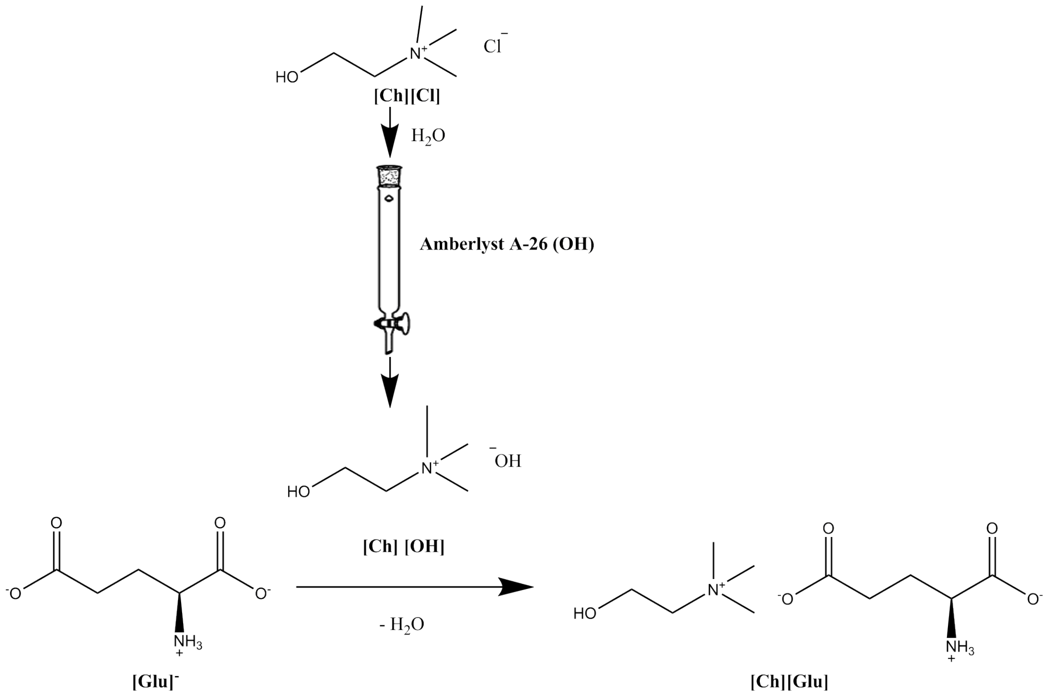

**Fig. S15: Schematic synthetic procedure for the preparation of [Ch][Glu] IL.**

### SI Tables

**Table S1: $\Delta G^{0'}{}_{u}$ (T) for drkN SH3 in water and ILs.**

| Temperature (K) | $\Delta G^{0'}{}_{u}$ (kcal mol$^{-1}$) Water | Buffer | 0.10 M [Ch][Glu] | 0.35 M [Ch][Glu] | 0.15 M [Bmim][dca] |
|---|---|---|---|---|---|
| **278.1** | -0.16 ± 0.01 | 0.11 ± 0.01 | 0.16 ± 0.01 | 0.70 ± 0.04 | -0.95 ± 0.06 |
| **283.1** | -0.030 ± 0.002 | 0.22 ± 0.01 | 0.26 ± 0.02 | 0.77 ± 0.05 | -0.94 ± 0.06 |
| **288.1** | 0.035 ± 0.002 | 0.26 ± 0.02 | 0.31 ± 0.02 | 0.82 ± 0.05 | -0.98 ± 0.06 |
| **293.2** | -0.0012 ± 0.0001 | 0.23 ± 0.01 | 0.30 ± 0.02 | 0.83 ± 0.05 | -1.04 ± 0.06 |
| **298.2** | -0.12 ± 0.01 | 0.12 ± 0.01 | 0.21 ± 0.01 | 0.75 ± 0.05 | -1.26 ± 0.08 |
| **303.2** | -0.30 ± 0.02 | -0.05 ± 0.003 | 0.028 ± 0.002 | 0.65 ± 0.04 | -1.5 ± 0.1 |
| **308.1** | -0.61 ± 0.04 | -0.34 ± 0.02 | -0.27 ± 0.02 | 0.41 ± 0.03 | -2.0 ± 0.1 |
| **313.1** | -1.12 ± 0.07 | -0.81 ± 0.05 | -0.72 ± 0.04 | 0.047 ± 0.003 | -2.7 ± 0.2 |

$\Delta G^{0'}{}_{u}$ ($T$) of drkN SH3 was calculated as described in the main text and methods section. The values are based on peak volume (simultaneously for folded and unfolded states) of indole side chain of W36 drkN SH3 in water, buffer 0.05 M sodium phosphate pH 7.2, 0.10 and 0.35 M [Ch][Glu], and 0.15 M [Bmim][dca], from 278 K to 313 K. Error represent the propagation of uncertainty in peak volume estimated by PINT. Details can be found in the SI Materials and Methods section.

**Table S2: Thermodynamic parameters for SH3 in the presence of ILs.**

| | $T_m$, K | $\Delta H^{0'}{}_{u}$ ($T_m$), kcal mol$^{-1}$ | $T_s$, K | $\Delta H^{0'}{}_{u}$ ($T_s$), kcal mol$^{-1}$ | $\Delta C_p{}^{0'}{}_{u}$ kcal mol$^{-1}$ K$^{-1}$ | $\Delta G^{0'}{}_{u}$ ($T_{m,water}$), kcal mol$^{-1}$ | $\Delta H^{0'}{}_{u}$ ($T_{m,water}$), kcal mol$^{-1}$ | $T\Delta S^{0'}{}_{u}$ ($T_{m,water}$), kcal mol$^{-1}$ |
|---|---|---|---|---|---|---|---|---|
| **Water** | **294 ± 1** | 6 ± 1 | **289.0 ± 0.5** | 0.06 ± 0.02 | 1.16 ± 0.07 | **0** | **6.2 ± 0.5** | **6.2 ± 0.5** |
| **Buffer (0.05 M Sodium Phosphate)** | **301.4 ± 0.4** | 13.6 ± 0.8 | **288.7 ± 0.6** | 0.29 ± 0.02 | 1.05 ± 0.07 | **0.23 ± 0.01** | **6.2 ± 0.2** | **6.0 ± 0.2** |
| **0.10 M [Ch][Glu]** | **303.0 ± 0.6** | 15.0 ± 0.8 | **289.1 ± 0.5** | 0.35 ± 0.02 | 1.06 ± 0.07 | **0.30 ± 0.02** | **5.9 ± 0.1** | **5.6 ± 0.1** |
| **0.35 M [Ch][Glu]** | **314.5 ± 0.6** | 21 ± 1 | **289.6 ± 0.7** | 0.86 ± 0.02 | 0.83 ± 0.07 | **0.83 ± 0.05** | **4.8 ± 0.1** | **4.0 ± 0.1** |
| **0.15 M [Bmim][dca]** | NA | NA | **285 ± 1** | -0.89 ± 0.03 | 1.3 ± 0.1 | **-1.08 ± 0.06** | **11 ± 2** | **12 ± 2** |

$\Delta G^{0'}{}_{u}$ ($T$) were calculated as described in the main text and methods section. $T_m$, $T_s$, $\Delta H^{0'}{}_{u}$ ($T_m$), $\Delta H^{0'}{}_{u}$ ($T_s$), and $\Delta C_p{}^{0'}{}_{u}$ were obtained by fitting **Eq. 2** when $T_m$ cannot be defined. Uncertainties represent the standard deviation from the fitting. The extrapolation of $\Delta H^{0'}{}_{u}$ ($T_{m,water}$) and $\Delta S^{0'}{}_{u}$ ($T_{m,water}$) and their uncertainties to $T_{m,water} \approx 294$ K ($T_m$ in cosolute-free solution), using $\Delta H^{0'}{}_{u}$ ($T_m$) or $\Delta H^{0'}{}_{u}$ ($T_s$) when $T_m$ cannot be defined, was obtained by Kirchhoff's temperature-dependent relations. See SI Materials and Methods for details. NA, not applicable.

**Table S3: Change in thermodynamic parameters for SH3 in the presence of ILs.**

| | $\Delta T_m$, K | $\Delta T_s$, K | $\Delta\Delta C_p{}^{0'}{}_u$, kcal mol$^{-1}$ | $\Delta\Delta G^{0'}{}_u$ ($T_{m,water}$), kcal mol$^{-1}$ | $\Delta\Delta H^{0'}{}_u$ ($T_{m,water}$), kcal mol$^{-1}$ | $T\Delta\Delta S^{0'}{}_u$ ($T_{m,water}$), kcal mol$^{-1}$ |
|---|---|---|---|---|---|---|
| **Buffer (0.05 M Sodium Phosphate)** | 7 ± 1 | -0.3 ± 0.7 | -0.1 ± 0.1 | **0.23 ± 0.01** | **0.0 ± 0.5** | **-0.2 ± 0.5** |
| **0.10 M [Ch][Glu]** | 9 ± 1 | 0.1 ± 0.7 | -0.1 ± 0.1 | **0.30 ± 0.02** | **-0.3 ± 0.5** | **-0.6 ± 0.5** |
| **0.35 M [Ch][Glu]** | 20 ± 1 | 0.6 ± 0.9 | -0.3 ± 0.1 | **0.83 ± 0.05** | **-1.4 ± 0.5** | **-2.2 ± 0.5** |
| **0.15 M [Bmim][dca]** | NA | -4 ± 1 | 0.1 ± 0.1 | **-1.04 ± 0.06** | **5 ± 2** | **6 ± 2** |

Excess changes are used as reference water for the thermodynamic parameters of drkN SH3. Uncertainties represent the error propagation from the initial standard fitting error. NA, not applicable.

**Table S4: Parameters for the drkN SH3 interconversion extracted from ZZex in water and aqueous-ILs.**

| | Water | | | | | | 0.35 M [Ch][Glu] IL | | | | | | 0.15 M [Bmim][dca] IL | | | | | |
|---|---|---|---|---|---|---|---|---|---|---|---|---|---|---|---|---|---|---|
| **Residue** | $p_f$ | $p_u$ | $R_{1f}$ ($s^{-1}$) | $R_{1u}$ ($s^{-1}$) | $k_u$ ($s^{-1}$) | $k_f$ ($s^{-1}$) | $p_f$ | $p_u$ | $R_{1f}$ ($s^{-1}$) | $R_{1u}$ ($s^{-1}$) | $k_u$ ($s^{-1}$) | $k_f$ ($s^{-1}$) | $p_f$ | $p_u$ | $R_{1f}$ ($s^{-1}$) | $R_{1u}$ ($s^{-1}$) | $k_u$ ($s^{-1}$) | $k_f$ ($s^{-1}$) |
| A3 | 0.55 | 0.45 | 2.50 | 2.16 | 0.44 | 0.54 | 0.85 | 0.15 | 2.02 | 2.10 | 0.13 | - | 0.20 | 0.80 | 4.04 | 2.38 | 0.27 | 0.07 |
| I4 | 0.51 | 0.49 | 2.50 | 2.00 | 0.86 | - | 0.82 | 0.18 | 2.08 | 2.85 | 0.14 | 0.64 | 0.10 | 0.90 | 1.30 | 1.84 | 1.55 | 0.18 |
| A5 | 0.51 | 0.49 | 2.69 | 2.27 | 0.59 | 0.61 | 0.82 | 0.18 | 2.29 | 2.68 | - | - | - | - | - | - | - | - |
| D8 | 0.59 | 0.41 | 3.36 | 2.34 | - | - | 0.88 | 0.12 | 1.93 | 0.50 | - | 0.64 | - | - | - | - | - | - |
| **S10** | **0.61** | **0.39** | **2.84** | **2.37** | **0.62** | **0.97** | **0.79** | **0.21** | **1.93** | **2.45** | **0.15** | **0.56** | **0.27** | **0.73** | **3.40** | **2.51** | **0.67** | **0.25** |
| **T12** | **0.45** | **0.55** | **2.36** | **2.46** | **0.78** | **0.63** | **0.88** | **0.12** | **1.88** | **2.00** | **0.15** | **1.12** | **0.16** | **0.84** | **2.69** | **2.39** | **1.20** | **0.22** |
| **A13** | **0.58** | **0.42** | **2.35** | **2.33** | **0.62** | **0.86** | **0.79** | **0.21** | **1.96** | **2.44** | **0.14** | **0.54** | **0.26** | **0.74** | **2.62** | **2.30** | **0.91** | **0.31** |
| D15 | - | - | - | - | - | - | 0.59 | 0.41 | 1.96 | 1.87 | 0.24 | - | 0.15 | 0.85 | 2.23 | 2.39 | 1.29 | |
| **E16** | **0.46** | **0.54** | **2.50** | **2.52** | **0.47** | **0.40** | **0.87** | **0.13** | **2.05** | **2.46** | **0.16** | **1.04** | **0.16** | **0.84** | **2.85** | **2.33** | **0.54** | **0.11** |
| **S18** | **0.57** | **0.43** | **2.47** | **2.55** | **0.63** | **0.85** | **0.72** | **0.28** | **2.39** | **2.46** | **0.40** | **1.04** | **0.24** | **0.76** | **2.65** | **2.47** | **1.05** | **0.33** |
| F19 | 0.72 | 0.28 | 3.11 | 2.08 | 0.51 | 1.31 | 0.68 | 0.32 | 2.62 | 2.38 | 0.36 | 0.79 | 0.27 | 0.73 | 4.20 | 3.16 | 0.93 | 0.34 |
| R20 | 0.63 | 0.37 | 2.73 | 1.83 | 0.57 | 0.97 | 0.49 | 0.51 | 2.50 | 2.52 | - | 0.29 | 0.10 | 0.90 | 3.31 | 2.51 | 0.62 | 0.07 |
| **T22** | **0.59** | **0.41** | **2.71** | **2.50** | **0.67** | **0.98** | **0.80** | **0.20** | **2.09** | **2.63** | **0.18** | **0.72** | **0.23** | **0.77** | **2.57** | **2.55** | **1.26** | **0.37** |
| **L25** | **0.63** | **0.37** | **2.65** | **1.43** | **0.53** | **0.91** | **0.92** | **0.08** | **2.03** | **1.95** | **0.10** | **1.24** | **0.17** | **0.83** | **2.02** | **2.22** | **1.16** | **0.24** |
| I27 | 0.53 | 0.47 | 2.39 | 2.14 | 0.77 | 0.87 | 0.64 | 0.36 | 1.82 | 1.88 | 0.20 | - | 0.14 | 0.86 | 1.70 | 2.00 | 1.04 | - |
| **L28** | **0.49** | **0.51** | **2.40** | **2.37** | **0.77** | **0.74** | **0.84** | **0.16** | **2.00** | **2.48** | **0.09** | **0.49** | **0.15** | **0.85** | **1.90** | **2.22** | **1.30** | **0.22** |
| **N29** | **0.66** | **0.34** | **2.60** | **2.07** | **0.54** | **1.07** | **0.82** | **0.18** | **2.06** | **1.51** | **0.13** | **0.58** | **0.30** | **0.70** | **2.60** | **2.33** | **0.98** | **0.42** |
| M30 | 0.47 | 0.53 | 2.76 | 2.68 | 1.05 | 0.94 | - | - | - | - | - | - | - | - | - | - | - | - |
| R38 | 0.55 | 0.45 | 2.50 | 2.51 | 0.49 | 0.60 | 0.84 | 0.16 | 2.00 | 2.48 | 0.09 | 0.49 | 0.18 | 0.82 | 2.57 | 2.37 | - | 0.17 |
| **A39** | **0.40** | **0.60** | **2.34** | **2.68** | **0.87** | **0.59** | **0.84** | **0.16** | **2.06** | **2.55** | **0.14** | **0.77** | **0.14** | **0.86** | **2.04** | **2.28** | **1.35** | **0.21** |
| E40 | 0.46 | 0.54 | 2.27 | 2.41 | 0.87 | | 0.81 | 0.19 | 1.92 | 2.18 | 0.18 | 0.80 | 0.15 | 0.85 | 2.59 | 2.32 | 0.72 | 0.13 |
| L41 | 0.60 | 0.40 | 2.58 | 3.10 | 0.57 | 0.84 | 0.85 | 0.15 | 1.94 | 1.94 | 0.18 | 1.06 | 0.18 | 0.82 | 2.33 | 2.22 | 0.84 | 0.18 |
| **D42** | **0.48** | **0.52** | **2.68** | **1.98** | **0.68** | **0.64** | **0.86** | **0.14** | **2.00** | **2.13** | **0.12** | **0.73** | **0.12** | **0.88** | **2.17** | **2.48** | **1.55** | **0.20** |
| G43 | 0.55 | 0.45 | 2.46 | 2.37 | 0.69 | 0.84 | 0.74 | 0.26 | 2.13 | 2.61 | 0.01 | 0.02 | 0.20 | 0.80 | 2.44 | 2.32 | 1.12 | 0.27 |
| **G46** | **0.56** | **0.44** | **2.56** | **2.28** | **0.63** | **0.80** | **0.86** | **0.14** | **2.00** | **2.13** | **0.12** | **0.73** | **0.21** | **0.79** | **3.07** | **2.45** | **0.90** | **0.24** |
| L47 | 0.37 | 0.63 | 2.42 | 2.57 | 0.54 | 0.31 | 0.74 | 0.26 | 2.13 | 2.61 | 0.01 | 0.02 | 0.12 | 0.88 | 2.07 | 2.21 | - | 0.16 |
| I48 | 0.40 | 0.60 | 2.20 | 2.21 | 0.91 | 0.62 | 0.74 | 0.26 | 2.01 | 2.27 | 0.19 | 0.52 | 0.10 | 0.90 | 1.53 | 2.01 | - | 0.15 |
| S50 | 0.61 | 0.39 | 2.79 | 2.40 | - | 1.03 | 0.88 | 0.12 | 2.09 | 1.98 | - | 1.18 | 0.26 | 0.74 | 2.69 | 2.50 | - | 0.41 |
| Y52 | 0.50 | 0.50 | 2.69 | 2.23 | 0.57 | 0.56 | 0.80 | 0.20 | 2.16 | 1.65 | 0.06 | 0.25 | 0.16 | 0.84 | 2.34 | 2.12 | 1.08 | - |
| **I53** | **0.50** | **0.50** | **2.55** | **2.30** | **0.75** | **0.75** | **0.82** | **0.18** | **2.12** | **2.31** | **0.17** | **0.75** | **0.14** | **0.86** | **1.98** | **2.22** | **1.18** | **0.20** |
| E54 | 0.42 | 0.58 | 2.40 | 2.52 | - | 0.44 | 0.76 | 0.24 | 2.05 | 2.77 | - | 0.36 | 0.12 | 0.88 | 2.61 | 2.29 | - | 0.10 |
| M55 | 0.51 | 0.49 | 2.81 | 2.57 | 0.49 | 0.50 | 0.85 | 0.15 | 2.19 | 2.01 | 0.10 | 0.54 | 0.17 | 0.83 | 3.88 | 2.63 | 0.52 | - |
| **K56** | **0.62** | **0.38** | **2.69** | **2.30** | **0.62** | **1.02** | **0.86** | **0.14** | **2.22** | **2.22** | **0.20** | **1.26** | **0.25** | **0.75** | **3.70** | **2.44** | **0.48** | **0.16** |
| **W36sc** | **0.50** | **0.50** | **2.24** | **2.08** | **0.49** | **0.49** | **0.85** | **0.15** | **2.19** | **2.01** | **0.10** | **0.54** | **0.16** | **0.84** | **2.48** | **1.93** | **0.77** | **0.15** |

Parameters were extracted by fitting the experimental data on both auto-peaks (*ff* and *uu*) and exchange cross peaks (*uf* and f*u*) simultaneously in MATLAB. Data extracted from a series of 2D ZZex spectra acquired at 293.2 K, 600.13 MHz with variable mixing times ranging from 10 to 750 ms. To avoid ambiguity, we use: $k_f \equiv k_{U\to F}$, $k_u \equiv k_{F\to U}$.

**Table S5: Rate constants of drkN SH3 interconversion.**

| Residue | Water | | 0.35 M [Ch][Glu] | | 0.15 M [Bmim[dca] | |
|---|---|---|---|---|---|---|
| | $k_u$ ($s^{-1}$) | $k_f$ ($s^{-1}$) | $k_u$ ($s^{-1}$) | $k_f$ ($s^{-1}$) | $k_u$ ($s^{-1}$) | $k_f$ ($s^{-1}$) |
| **S10** | 0.62 | 0.97 | 0.15 | 0.56 | 0.67 | 0.25 |
| | | | 0.21 [a] | 0.77 [a] | 0.68 [a] | 0.26 [a] |
| **T12** | 0.78 | 0.63 | 0.15 | 1.12 | 1.20 | 0.22 |
| | | | 0.21 [a] | 1.56 [a] | 1.22 [a] | 0.23 [a] |
| **A13** | 0.62 | 0.86 | 0.14 | 0.54 | 0.91 | 0.31 |
| | | | 0.20 [a] | 0.75 [a] | 0.92 [a] | 0.32 [a] |
| **E16** | 0.47 | 0.40 | 0.16 | 1.04 | 0.54 | 0.11 |
| | | | 0.22 [a] | 1.45 [a] | 0.55 [a] | 0.11 [a] |
| **S18** | 0.63 | 0.85 | 0.40 | 1.04 | 1.05 | 0.33 |
| | | | 0.55 [a] | 1.45 [a] | 1.06 [a] | 0.33 [a] |
| **T22** | 0.67 | 0.98 | 0.18 | 0.72 | 1.26 | 0.37 |
| | | | 0.25 [a] | 0.99 [a] | 1.28 [a] | 0.38 [a] |
| **L25** | 0.53 | 0.91 | 0.10 | 1.24 | 1.16 | 0.24 |
| | | | 0.14 [a] | 1.72 [a] | 1.18 [a] | 0.25 [a] |
| **L28** | 0.77 | 0.74 | 0.09 | 0.49 | 1.30 | 0.22 |
| | | | 0.13 [a] | 0.68 [a] | 1.32 [a] | 0.22 [a] |
| **N29** | 0.54 | 1.07 | 0.13 | 0.58 | 0.98 | 0.42 |
| | | | 0.18 [a] | 0.80 [a] | 1.00 [a] | 0.43 [a] |
| **A39** | 0.87 | 0.59 | 0.14 | 0.77 | 1.35 | 0.21 |
| | | | 0.20 [a] | 1.07 [a] | 1.37 [a] | 0.22 [a] |
| **D42** | 0.68 | 0.64 | 0.12 | 0.73 | 1.55 | 0.20 |
| | | | 0.17 [a] | 1.01 [a] | 1.57 [a] | 0.21 [a] |
| **G46** | 0.63 | 0.80 | 0.12 | 0.73 | 0.90 | 0.24 |
| | | | 0.17 [a] | 1.01 [a] | 0.91 [a] | 0.25 [a] |
| **I53** | 0.75 | 0.75 | 0.17 | 0.75 | 1.18 | 0.20 |
| | | | 0.24 [a] | 1.04 [a] | 1.20 [a] | 0.20 [a] |
| **K56** | 0.62 | 1.02 | 0.20 | 1.26 | 0.48 | 0.16 |
| | | | 0.28 [a] | 1.75 [a] | 0.49 [a] | 0.16 [a] |
| **W36sc** | 0.49 | 0.49 | 0.10 | 0.54 | 0.77 | 0.15 |
| | | | 0.13 | 0.75 [a] | 0.78 [a] | 0.15 [a] |
| **Average** | **0.6 ± 0.1** | **0.8 ± 0.2** | **0.16 ± 0.07** | **0.8 ± 0.3** | **1.0 ± 0.3** | **0.24 ± 0.09** |
| | | | **0.2 ± 0.1[a]** | **1.1 ± 0.4[a]** | **1.0 ± 0.3[a]** | **0.25 ± 0.09[a]** |

Rate constants of accurately followed residues by ZZex experiments for water and aqueous-ILs conditions (0.35 M [Ch][Glu] and 0.15 M [Bmim][dca]) at 293.2 K, 600.13 MHz. Uncertainties represent the standard deviation from the average. [a] Viscosity-corrected rate. The relative viscosity ($\eta_{rel}$) used in 0.35 M [ChGlu] and 0.15 M [Bmim][dca] is 1.39 and 1.02, respectively.

**Table S6: Activation parameters for SH3 folding and unfolding.**

| | water | | | 0.35 M [Ch][Glu] IL | | | 0.15 M [Bmim][dca] IL | | |
|---|---|---|---|---|---|---|---|---|---|
| | **ΔG F→ TS‡ kcal mol$^{-1}$** | **ΔG U→TS‡ kcal mol$^{-1}$** | **$\Delta G_u$ kcal mol$^{-1}$** | **ΔG F→ TS‡ kcal mol$^{-1}$** | **ΔG U→TS‡ kcal mol$^{-1}$** | **$\Delta G_u$ kcal mol$^{-1}$** | **ΔG F→ TS‡ kcal mol$^{-1}$** | **ΔG U→TS‡ kcal mol$^{-1}$** | **$\Delta G_u$ kcal mol$^{-1}$** |
| **S10** | 17.43 | 17.17 | 0.26 | 18.25<br>18.06 [a] | 17.49<br>17.30 [a] | 0.76 | 17.38<br>17.37 [a] | 17.95<br>17.94 [a] | -0.57 |
| **T12** | 17.29 | 17.42 | -0.12 | 18.24<br>18.05 [a] | 17.08<br>16.89 [a] | 1.16 | 17.04<br>17.03 [a] | 18.04<br>18.02 [a] | -0.98 |
| **A13** | 17.43 | 17.23 | 0.19 | 18.29<br>18.10 [a] | 17.51<br>17.32 [a] | 0.78 | 17.21<br>17.20 [a] | 17.83<br>17.82 [a] | -0.62 |
| **E16** | 17.59 | 17.69 | -0.10 | 18.21<br>18.02 [a] | 17.12<br>16.93 [a] | 1.09 | 17.51<br>17.50 [a] | 18.46<br>18.45 [a] | -0.95 |
| **S18** | 17.42 | 17.25 | 0.17 | 17.68<br>17.49 [a] | 17.13<br>16.94 [a] | 0.56 | 17.12<br>17.11 [a] | 17.80<br>17.79 [a] | -0.68 |
| **T22** | 17.38 | 17.16 | 0.22 | 18.16<br>17.97 [a] | 17.34<br>17.15 [a] | 0.82 | 17.01<br>17.01 [a] | 17.72<br>17.71 [a] | -0.71 |
| **L25** | 17.52 | 17.21 | 0.31 | 18.47<br>18.28 [a] | 17.02<br>16.83 [a] | 1.45 | 17.06<br>17.05 [a] | 17.97<br>17.97 [a] | -0.91 |
| **L28** | 17.30 | 17.33 | -0.03 | 18.55<br>18.36 [a] | 17.56<br>17.37 [a] | 0.99 | 17.00<br>16.99 [a] | 18.03<br>18.02 [a] | -1.03 |
| **N29** | 17.51 | 17.11 | 0.40 | 18.35<br>18.16 [a] | 17.47<br>17.28 [a] | 0.88 | 17.16<br>17.15 [a] | 17.65<br>17.64 [a] | -0.49 |
| **A39** | 17.23 | 17.46 | -0.23 | 18.28<br>18.09 [a] | 17.30<br>17.11 [a] | 0.98 | 16.98<br>16.97 [a] | 18.05<br>18.04 [a] | -1.08 |
| **D42** | 17.37 | 17.41 | -0.04 | 18.37<br>18.18 [a] | 17.33<br>17.14 [a] | 1.03 | 16.90<br>16.89 [a] | 18.08<br>18.07 [a] | -1.18 |
| **G46** | 17.42 | 17.28 | 0.14 | 18.37<br>18.18 [a] | 17.33<br>17.14 [a] | 1.03 | 17.21<br>17.20 [a] | 17.97<br>17.96 [a] | -0.76 |
| **I53** | 17.32 | 17.32 | 0.00 | 18.18<br>17.99 [a] | 17.32<br>17.13 [a] | 0.87 | 17.05<br>17.04 [a] | 18.09<br>18.08 [a] | -1.04 |
| **K56** | 17.43 | 17.14 | 0.29 | 18.09<br>17.90 [a] | 17.02<br>16.83 [a] | 1.07 | 17.57<br>17.56 [a] | 18.21<br>18.20 [a] | -0.64 |
| **W36sc** | 17.57 | 17.57 | 0.00 | 18.52<br>18.33 [a] | 17.51<br>17.32 [a] | 1.01 | 17.30<br>17.29 [a] | 18.26<br>18.25 [a] | -0.95 |
| **Average** | 17.4 ± 0.1 | 17.3 ± 0.2 | 0.1 ± 0.2 | 18.3 ± 0.2<br>18.1 ± 0.2 [a] | 17.3 ± 0.2<br>17.1 ± 0.2 [a] | 1.0 ± 0.2 | 17.2 ± 0.2<br>17.2 ± 0.2 [a] | 18.0 ± 0.2<br>18.0 ± 0.2 [a] | -0.8 ± 0.2 |

Values per residue calculated as described in the main text and methods section. $\Delta G_{F\rightarrow TS‡}$ and $\Delta G_{U\rightarrow TS‡}$ are the modified standard-state activation free energies for unfolding and folding, respectively. Uncertainties represent the standard deviation from the average. [a]Viscosity-corrected rate. The relative viscosity ($\eta_{rel}$) used in 0.35 M [ChGlu] and 0.15 M [Bmim][dca] is 1.39 and 1.02, respectively.

**Table S7: Activation parameters and excess changes for SH3 folding and unfolding.**

| | $\Delta G^{0'\ddagger}_{F\rightarrow TS\ddagger}$, kcal mol$^{-1}$ | $\Delta G^{0'\ddagger}_{U\rightarrow TS\ddagger}$, kcal mol$^{-1}$ | $\Delta G^{0'}_{u}$, kcal mol$^{-1}$ | $\Delta\Delta G^{0'\ddagger}_{F\rightarrow TS\ddagger}$, kcal mol$^{-1}$ | $\Delta\Delta G^{0'\ddagger}_{U\rightarrow TS\ddagger}$, kcal mol$^{-1}$ | $\Delta\Delta G^{0'}_{u}$, kcal mol$^{-1}$ | α, $\Delta\Delta G^{0'\ddagger}_{U\rightarrow TS\ddagger}/\Delta\Delta G^{0'}_{u}$, kcal mol$^{-1}$ |
|---|---|---|---|---|---|---|---|
| **Water** | 17.4 ± 0.1 | 17.3 ± 0.2 | 0.1 ± 0.2 | **-** | **-** | - | - |
| **0.35 M [Ch][Glu]** | 18.3 ± 0.2 | 17.3 ± 0.2 | 1.0 ± 0.2 | **0.9 ± 0.2** | **0.0 ± 0.3** | **0.9 ± 0.3** | **0.2 ± 0.6** [a] |
| | 18.1 ± 0.2 [a] | 17.1 ± 0.2 [a] | | **0.7 ± 0.2** [a] | **-0.2 ± 0.3** [a] | | |
| **0.15 M [Bmim][dca]** | 17.2 ± 0.2 | 18.0 ± 0.2 | -0.8 ± 0.2 | **-0.2 ± 0.2** | **0.7 ± 0.1** | **-0.9 ± 0.1** | **0.7 ± 0.3** [a] |
| | 17.2 ± 0.2 [a] | 18.0 ± 0.2 [a] | | **-0.3 ± 0.2** [a] | **0.7 ± 0.1** [a] | | |

Average 15-residue values calculated as described in the main text and excess changes using water as a reference. Uncertainties represent the standard deviation from the average. [a] Viscosity-corrected rate.

**Table S8: Measured viscosities for aqueous-[Ch][Glu] solutions.**

| | Aqueous-[Ch][Glu] viscosity (cP) | | | |
|---|---|---|---|---|
| **Temperature (K)** | **0.1 M** | **0.5 M** | **1.0 M** | **1.5 M** |
| **298.2** | 1.113 ± 0.006 | 1.417 ± 0.005 | 2.04 ± 0.01 | 3.26 ± 0.06 |
| **304.2** | 0.993 ± 0.007 | 1.26 ± 0.01 | 1.797 ± 0.009 | 2.82 ± 0.03 |
| **310.2** | 0.89 ± 0.01 | 1.13 ± 0.02 | 1.598 ± 0.006 | 2.47 ± 0.02 |

The uncertainties are the standard deviation from the triplicate dataset.
